# Novel autoregulatory cases of alternative splicing coupled with nonsense-mediated mRNA decay

**DOI:** 10.1101/464404

**Authors:** Dmitri Pervouchine, Yaroslav Popov, Andy Berry, Beatrice Borsari, Adam Frankish, Roderic Guigó

## Abstract

Nonsense-mediated decay (NMD) is a eukaryotic mRNA surveillance system that selectively degrades transcripts with premature termination codons (PTC). Many RNA-binding proteins (RBP) regulate their expression levels by a negative feedback loop, in which RBP binds its own pre-mRNA and causes alternative splicing to introduce a PTC. We present a bioinformatic framework to identify novel such autoregulatory feedback loops by combining eCLIP assays for a large panel of RBPs with the data on shRNA inactivation of NMD pathway, and shRNA-depletion of RBPs followed by RNA-seq. We show that RBPs frequently bind their own pre-mRNAs and respond prominently to NMD pathway disruption. Poison and essential exons, i.e., exons that trigger NMD when included in the mRNA or skipped, respectively, respond oppositely to the inactivation of NMD pathway and to the depletion of their host genes, which allows identification of novel autoregulatory mechanisms for a number of human RBPs. For example, SRSF7 binds its own pre-mRNA and facilitates the inclusion of two poison exons; SFPQ binding promotes switching to an alternative distal 3’-UTR that is targeted by NMD; RPS3 activates a poison 5’-splice site in its pre-mRNA that leads to a frame shift; U2AF1 binding activates one of its two mutually exclusive exons, leading to NMD; TBRG4 is regulated by cluster splicing of its two essential exons. Our results indicate that autoregulatory negative feedback loop of alternative splicing and NMD is a generic form of post-transcriptional control of gene expression.

## Introduction

Gene expression in higher eukaryotes is regulated at many different levels. The output of the transcriptional program is maintained by a large number of protein factors and cis-regulatory elements, which control the balance between mRNA production and degradation [1, 2]. Nonsense mutations and frame-shifting splicing errors induce premature termination codons (PTC) that give rise to mRNAs encoding truncated, dysfunctional proteins. In eukaryotic cells, mRNA transcripts with PTC are selectively degraded by the surveillance mechanism called Nonsense-Mediated mRNA Decay (NMD) [3].

The so-called exon junction complex-dependent (EJC) model postulates that NMD distinguishes between normal and premature translation termination in the cytoplasm, where ribosomes displace EJCs from within, but not downstream of the reading frame [4, 5]. These complexes are deposited approximately 50 nucleotides (nt) upstream of the exon-exon junctions during pre-mRNA splicing. EJCs that remain associated with the mRNA after the initial round of translation serve as indicators of whether the termination codon is premature or not, because the normal termination codons are usually located in the last exon. The presence of EJCs downstream of the stop codon triggers a cascade of events, in which the up-frameshift 1 factor (UPF1) plays a central role [5]. The phosphorylated UPF1 recruits the endonuclease SMG6 and other factors causing deadenylation and decapping, targeting the cleaved mRNA for degradation by cellular exonucleases [5]. Other models propose that sensing the distinction between a normal termination codon and a PTC depends on the distance between the terminating ribosome and the poly(A) tail, in which the interaction of eRF3 with PABPC1 is important, or that an early ribosome release caused by the PTC exposes the downstream unprotected mRNA to degradation by nucleases independently of EJCs [6].

It has been increasingly reported over past years that NMD is not only dedicated to the destruction of PTC-containing mRNAs that appear as a result of nonsense mutations or splicing errors, but that it also plays a key role in regulating the expression of a broad class of physiological transcripts [7, 8]. Targets of NMD include tissue-specific transcripts [9], transcripts with mutually exclusive exons [10], mRNAs with upstream open reading frames (uORFs) and long 3’-untranslated regions (UTRs) [11], and transcripts emanating from transposons and retroviruses [12]. The mechanism, in which the cell employs alternative splicing (AS) coupled with NMD to downregulate the abundance of mRNA transcripts, shortly termed AS-NMD [13] (also referred to as regulated unproductive splicing and translation [7] or unproductive splicing [14]), is found in all eukaryotes that have been studied to date and often exhibits a high degree of evolutionary conservation [15, 16]. Current records in GENCODE database indicate that up to one-third of human protein-coding genes have at least one annotated transcript that would be degraded by NMD, and that a significant fraction of them have mammalian orthologs, suggesting that unproductive splicing is a widespread and functionally selected mechanism of post-transcriptional control of gene expression [17].

NMD is involved in the development of cancers, where it can downregulate the expression of tumor-suppressor genes or activate the expression of oncogenes that are normally suppressed [18]. In several cancers including hepatocellular carcinoma, alternative splicing of the Kruppel-like factor 6 (KLF6) tumor suppressor gene leads to a pathogenic AS-NMD splice variant associated with increased tumor metastasis and mortality [19]. Inhibition of the NMD pathway stabilizes many transcripts necessary for tumorigenesis [20]. Intriguingly, a substantial number of long non-coding RNAs and snoRNAs are also found to be substrates of NMD, suggesting that NMD pathway is not exclusively dedicated to mRNAs [21–23]. Among them is the metastasis-associated lung adenocarcinoma transcript 1 (MALAT1), which is downregulated in gastric cancer upon UPF1 overexpression [24].

Many RNA-binding proteins (RBPs), and particularly splicing factors (SFs), control their own expression levels by a negative feedback loop mediated by AS-NMD, in which the excessive amount of RBP binds its own pre-mRNA and causes alternative splicing to induce a PTC. Studies have shown that mutations in RBP binding sites abolish this negative feedback, while RBP overexpression leads to the increased fraction of unproductively-spliced mRNA [25]. To date, many genes that utilize this mechanism are known, including SR proteins [14, 26, 27], hnRNP family members (28–30), TDP-43 [31, 32], TRA2β [33], MBNL [34, 35], PTB [36], CHTOP [37], FUS [38], and core spliceosomal and ribosomal proteins (39–41). Besides autoregulation, some RBPs use AS-NMD to cross-regulate the expression of their family members. Examples include hnRNPL/hnRNPLL [30], PTBP1/PTBP2 [36, 42], RBM10/ RBM5 [39], RBFOX2/PTBP2 [43], RBFOX2/RBFOX3 [44] and others (see [45] for the detailed analysis of cross-regulatory networks). Disruption of auto- or cross-regulation of SFs is associated with human pathogenic states, including neurodegenerative diseases and cancer [39, 46–48].

The connection between AS event and the position of PTC that is induced by it is not always evident because PTC may appear anywhere downstream of the frameshift. The simplest and the most studied case is the so-called poison cassette exon, i.e. cassette exon with an early in-frame PTC, which is normally skipped, but triggers degradation by NMD when included in the mature mRNA [14] (Figure 1a). The classic examples of auto- and cross-regulatory poison exons are documented in serine/arginine-rich (SR) proteins [14]. However, other classes of AS events such as alternative 5’- and 3’-splice sites or intron retention also contribute to AS-NMD [49]. A case that is reciprocal to poison exons, termed here as “essential” exon, occurs when a cassette exon, which is normally included in the mature mRNA, triggers NMD when skipped. For instance, alternative skipping of PTB exon 11 yields an mRNA that is removed by NMD, and the skipping is itself promoted by PTB in a negative feedback loop [36]. Similarly, the spliceosomal RNA binding protein RBM10, which is associated with TARP syndrome and lung adenocarcinoma, downregulates its own expression and that of RBM5 by promoting skipping of several its essential exons [39].

**Figure 1.**
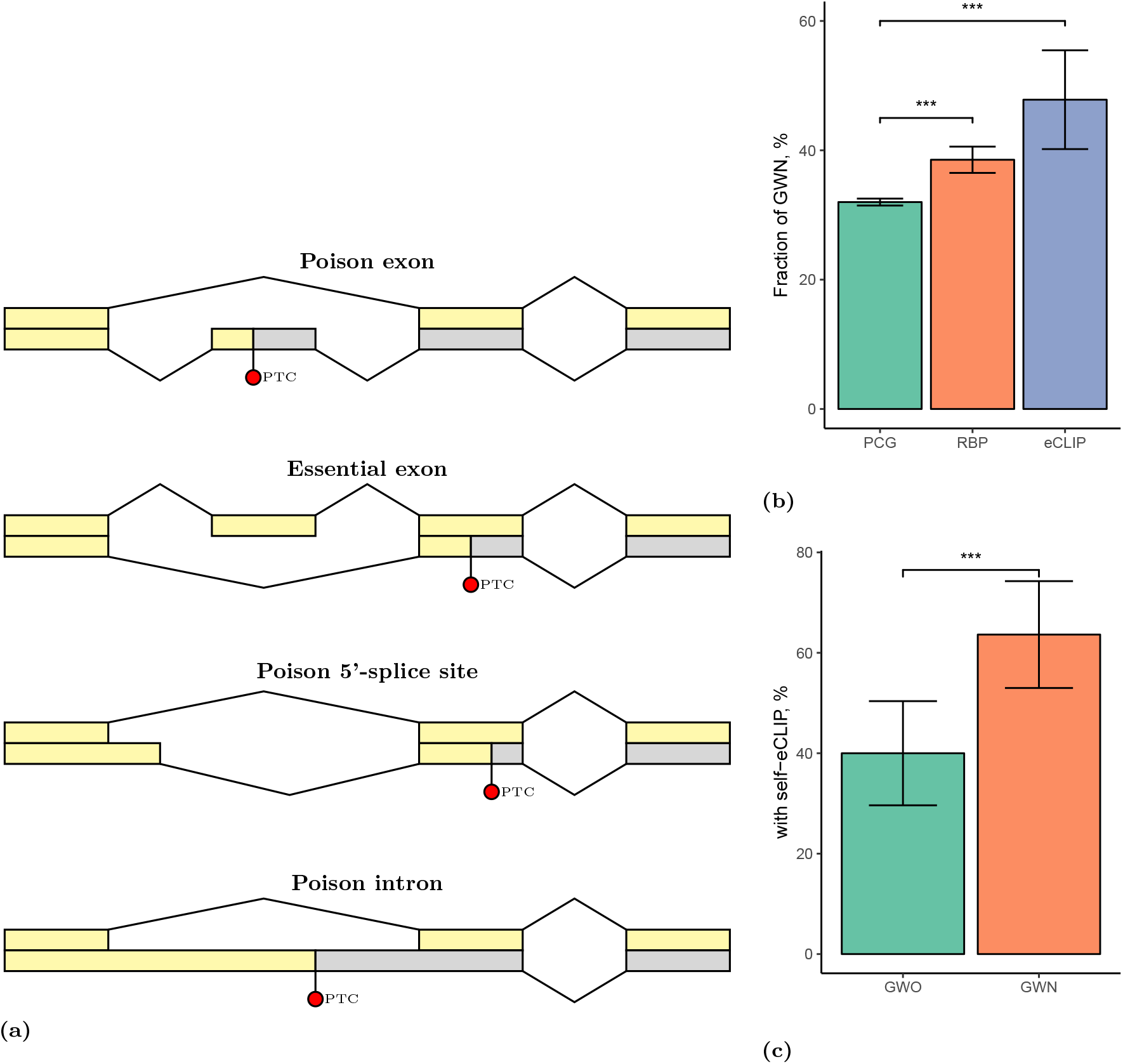
**(a)** The relationship between an alternative splicing event and the PTC that is induced by it. Top-down: a poison exon causes NMD when included; an essential exon causes NMD when skipped; a poison 5’-splice site causes a frame shift and induces a downstream PTC; a retained poison intron carries a PTC. **(b)** The proportion of genes with NMD (GWN) among all protein coding genes (PCG), RNA-binding proteins (RBP), and RNA-binding proteins that were profiled by eCLIP (eCLIP). (**c**) The proportion of RBPs that bind their own pre-mRNA among genes with NMD (GWN) and genes without NMD (GWO). Significant differences in proportions are shown by asterisks.

In this work, we approach the question of finding novel autoregulatory AS-NMD feedback loops among RBPs. We combine three publicly available data sources: the enhanced crosslinking and immunoprecipitation (eCLIP) assay for a large panel of RBPs [50], short-hairpin RNA (shRNA) knockdowns (KD) of the same RBPs followed by RNA sequencing (RNA-seq) [51–53], and a model of NMD pathway inactivation by shRNA co-depletion of UPF1 and XRN1 followed by RNA-seq [54]. In particular, we show that (i) RBPs with annotated NMD transcripts tend to bind their own pre-mRNAs more frequently than do other RBPs; (ii) RBPs are enriched among genes that respond to the inactivation of NMD pathway; (iii) poison and essential exons react oppositely to the disruption of NMD pathway and, in the case of a negative AS-NMD feedback loop, to perturbations of the expression level of their host genes. Based on this logic, we identified a set of exons that significantly and substantially change their inclusion level in response to the inactivation of NMD pathway components and to downregulation of the host gene. We analyzed these exons together with eCLIP data and proposed a number of AS-NMD autoregulatory mechanisms for human RBPs, including serine/arginine-rich and proline/glutamine-rich splicing factors SRSF7 and SFPQ, human ribosomal protein RPS3, spliceosomal auxiliary factor U2AF1, and Fas-activated serine/threonine kinase domain-containing protein TBRG4. These predicted autoregulatory mechanisms are outlined in last section.

## Materials and Methods

### Genomes and transcript annotations

February 2009 assembly of the human genome (hg19, GRCh37) was downloaded from Genome Reference Consortium [55]. The respective transcript annotation v19 was downloaded from GENCODE website [56]. Transcript annotations were parsed by custom scripts to extract positions of introns and exons. Intron and exon sequences were extracted using bedtools getfasta tool [57]. A transcript was considered as NMD target if it was labelled as “nonsense_mediated_decay” by GENCODE. A human protein-coding gene will be referred to as Gene With NMD (GWN) if it contains at least one transcript isoform annotated as NMD [17], and Gene Without NMD (GWO) otherwise.

The annotated exon [*x, y*] with the acceptor site x and the donor site y is defined to be a cassette exon if there exist introns [*a,x*] and [*y,b*] such that [*a,b*] is also an intron, i.e. the intervals [*a,x*], [*y,b*], and [*a,b*] are introns in at least one annotated transcript. A cassette exon is defined to be a poison exon, if it contains a stop codon of an annotated NMD-transcript. For essential exons, we use a different definition to avoid connecting exon skipping to the induced downstream PTC. For each exon in each annotated CDS, we check whether it is essential or not by removing its nucleotide sequence from the transcript, translating the modified nucleotide sequence to aminoacids, and checking if a PTC appears 50 nt upstream of at least one splice junction. A relative position of an exon in a transcript was defined as the position of its midpoint normalized by a linear transformation to the range from 0% to 100%, where the 5’-exon is 0% and the 3’-exon is 100%.

### RNA-binding proteins (RBP)

The list of genes with an annotated RNA-binding function was obtained by searching for the term “RNA binding” (GO:0003723) in the table that was obtained by merging ENSEMBL identifiers of human genes with Gene Ontology annotation on UniProt identifiers [58, 59]. The resulting list of RBPs containing 1544 genes is shown in Supplementary Data File 1.

### Enhanced crosslinking and immunoprecipitation assay (eCLIP)

We used publicly available eCLIP data for 115 RBPs profiled in [50]. eCLIP peaks, which were called by the data producers, were downloaded from ENCODE data repository in bed format [52, 53]. The peaks in two immortalized human cell lines, K562 and HepG2, were filtered by the condition log *FC* ≥ 3 and P-value< 0.001 as recommended [50]. Since the agreement between peaks in the two replicates was moderate (the median Jaccard distance 25% and 28% in K562 and HepG2, respectively), we took the union of peaks between the two replicates within each cell line, and then pooled the resulting peaks between cell lines. A summary of eCLIP profiles that were used in this study and their accession numbers is given in Supplementary Table S1.

### Short-hairpin RNA knockdown of RBP followed by RNA-seq

Publicly available data on short-hairpin (shRNA) knockdown of 250 RBPs followed by RNA-seq (shRNA-KD) [51] were downloaded in BAM format from ENCODE data repository [52, 53]. A summary of RBP depletion data and the respective accession numbers is given in Supplementary Table S1. Exon inclusion metrics (PSI) were called for all annotated exons using IPSA software with the default settings [60]. PSI values of individual exons were averaged between bioreplicates.

### UPF1/XRN1 co-depletion analysis

We used the expression profiling by RNA-sequencing in HEK293 Flp-In T-Rex cells that were subjected to siRNA-mediated depletion of XRN1 and co-depletion of UPF1 (GEO accession GSE57433) [54]. Transcript quantification for target dataset (GSM1382448) vs. control (GSM1382445) were done using cufflinks2 by data producers [61]. The resulting read counts were processed in R statistics software by DESeq2 package using normal shrinkage correction [62]. Due to the fact that the original data was not replicated, the corrected p-values were not computed. Exon inclusion metrics (PSI) were called for all annotated exons using IPSA software with the default settings [60].

### Exon inclusion rate (PSI)

Genomic alignments of short reads from all RNA-seq experiments were processed using IPSA pipeline to obtain read counts supporting splice junctions [60]. Read counts were filtered by the entropy content of the offset distribution, annotation status and canonical GT/AG dinucleotides at splice sites [60]. The exon inclusion rate (ψ, Percent-Spliced-In, or PSI ratio) was calculated for exons of annotated transcripts according to the equation

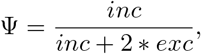

where *inc* is the corrected number of reads supporting exon inclusion and exc is the corrected number of reads supporting exon exclusion. Ψ values with the denominator below 20 counts were considered unreliable and discarded. This definition of exon inclusion rate applies not only to cassette exons, but also to other types of AS events, e.g., alternative 5’- and 3’-splice sites, whenever inclusion and exclusion reads allow successful discrimination between alternatively-spliced transcript isoforms.

The changes in exon inclusion were assessed by using ΔΨ = Ψ(*KD*) – Ψ(*Control*) metric, where Ψ(*KD*) and Ψ(*Control*) are exon inclusion rates in the KD experiment and in the control, respectively. The distribution of ΔΨ values spans the interval from −1 to 1 and, as in the case of gene expression, its mean and standard deviation depend on the level of gene expression. We used the number of reads in the denominator of Ψ as a proxy for the local gene expression level at a given exon, and developed the following heuristic procedure to assess statistical significance of exon inclusion changes.

We binned Ψ values of all exons by log_10_ of the mean split read count, which is the average of Ψ denominators between KD and the control. In each bin, we computed the mean and the standard deviation of ΔΨ, excluding exons with ΔΨ = 0, and assigned the corresponding *z*-score to each exon. The distribution of *z*-scores was bell-shaped and it was not unreasonable to assign a normal one-tail probability to each exon. The probability was then corrected by Šidák correction with the number of tests equal to the number of exons.

### Gene ontology analysis

We used Human Gene Ontology annotation provided by Gene Ontology (GO) Consortium [58, 63]. Enrichment of GO terms in gene sets of interest was done using GOstats library [64] within Bioconductor R package, and also using GOrilla, a tool for GO term enrichment analysis in ranked gene lists [65].

### Statistical analysis

The data were analyzed and visualized using R statistics software version 3.4.1 and `ggplot2` package. Tests for proportions were performed using normal approximation to binomial distribution for samples of size *n* > 40 without continuity correction. Non-parametric tests were performed by built-in R functions using normal approximation with continuity correction. Error bars in all figures correspond to the 95% confidence intervals. Wilcoxon one-sample test was used to assess ΔΨ distribution for departures from zero. One-sided P-values are reported throughout the paper.

## Results

### RBP often undergo NMD and frequently bind their own pre-mRNA

Throughout this paper, a human protein-coding gene with at least one annotated NMD transcript is referred to as GWN (Gene With NMD); otherwise it is referred to as GWO. First, we assessed the functional attribution of GWN set by Gene Ontology analysis [65]. Genes with annotated molecular function of RNA-binding, nucleotide and ribonucleotide binding, and genes involved in biological processes related to splicing were significantly enriched among GWN compared to GWO (Supplementary Table S2). In what follows, we focused on the analysis of genes that are both GWN and RBP.

Approximately one third of the annotated human protein-coding genes are GWN (32%, or 6,476 out of 20,242) [17]. We asked whether the proportion of GWN among specific gene classes is greater than that among all protein-coding genes (Figure 1b). Indeed, 595 out of 1,544 RBPs are GWN, which is significantly greater than the background proportion (38% vs. 32%, one-sample proportion test, *P* = 6· 10^−4^). Furthermore, 55 out 115 splicing-related RBPs that were profiled by eCLIP are GWN, indicating a greater GWN enrichment (48% vs. 32%, one-sample proportion test, *P* = 3 · 10^−4^). Thus, we confirm that RBP, and particularly splicing-related RBPs, as a class of genes have a higher propensity to undergo NMD than do other protein-coding genes.

We next asked whether RBPs tend to bind their own pre-mRNAs. To test this, we analyzed eCLIP profiles of 115 RBPs and intersected them with the genomic ranges that encode their cognate genes. While 35 out of 55 GWNs profiled by eCLIP contain at least one eCLIP peak in their own gene, the respective proportion for GWO is 24 out of 60, i.e., for GWN the proportion is significantly greater (64% vs. 40%, 2-sample proportion test, *P* = 0.005, Figure 1c). This enrichment is not due to an imbalance in eCLIP signal density since the number of eCLIP peaks that fall within RBP genes is not significantly different between GWN and GWO (Wilcoxon test, *P* = 0.15). This indicates that RBPs with annotated NMD events bind their own pre-mRNA more frequently than do RBPs without annotated NMD events. According to this statistical evidence, one may expect that RBPs frequently autoregulate their expression via AS-NMD. It is the aim of this paper to identify such cases.

### RBPs are enriched among UPF1/XRN1 co-depletion targets

The degradation of transcripts by NMD is initiated by UPF1-activated endonucleolytic cleavage of the nonsense RNA in the vicinity of the PTC followed by a rapid digestion by cytoplasmic 5’-3’ exonucle-ases [66, 67]. Specifically, the exonuclease XRN1 degrades the 3’-fragment derived from the endonucleolytic cleavage, as well as the decapped full-length nonsense RNA [66]. We analyzed publicly available data on transcriptome-wide identification of NMD substrates by UPF1/XRN1 co-depletion [54] and performed differential gene expression analysis. To quantify splicing changes, we computed ΔΨ metric for all exons of annotated transcripts as explained in Materials and Methods and assumed that exon inclusion levels change independently of gene expression levels.

The ontology analysis of genes that are substantially upregulated in UPF1/XRN1 co-depletion reveals a significant enrichment of genes related to splicing, including components of the heterogeneous nuclear ribonucleoprotein complexes (hnRNPs) and RNA-binding factors known to co-localize with core spliceosomal proteins (Figure 2a). The top 15 genes with the largest fold change in UPF1/XRN1 co-depletion and the respective GO terms are listed in Supplementary Tables S3 and S4, respectively. Although the lack of replication in the co-depletion experiment does not permit rigorous assignment of statistical significance, it can be noted that more genes are upregulated upon UPF1/XRN1 co-depletion compared to genes that are downregulated at the same value of *logFC*, indicating that the upregulated gene set contains natural NMD targets (Figure 2a).

**Figure 2.**
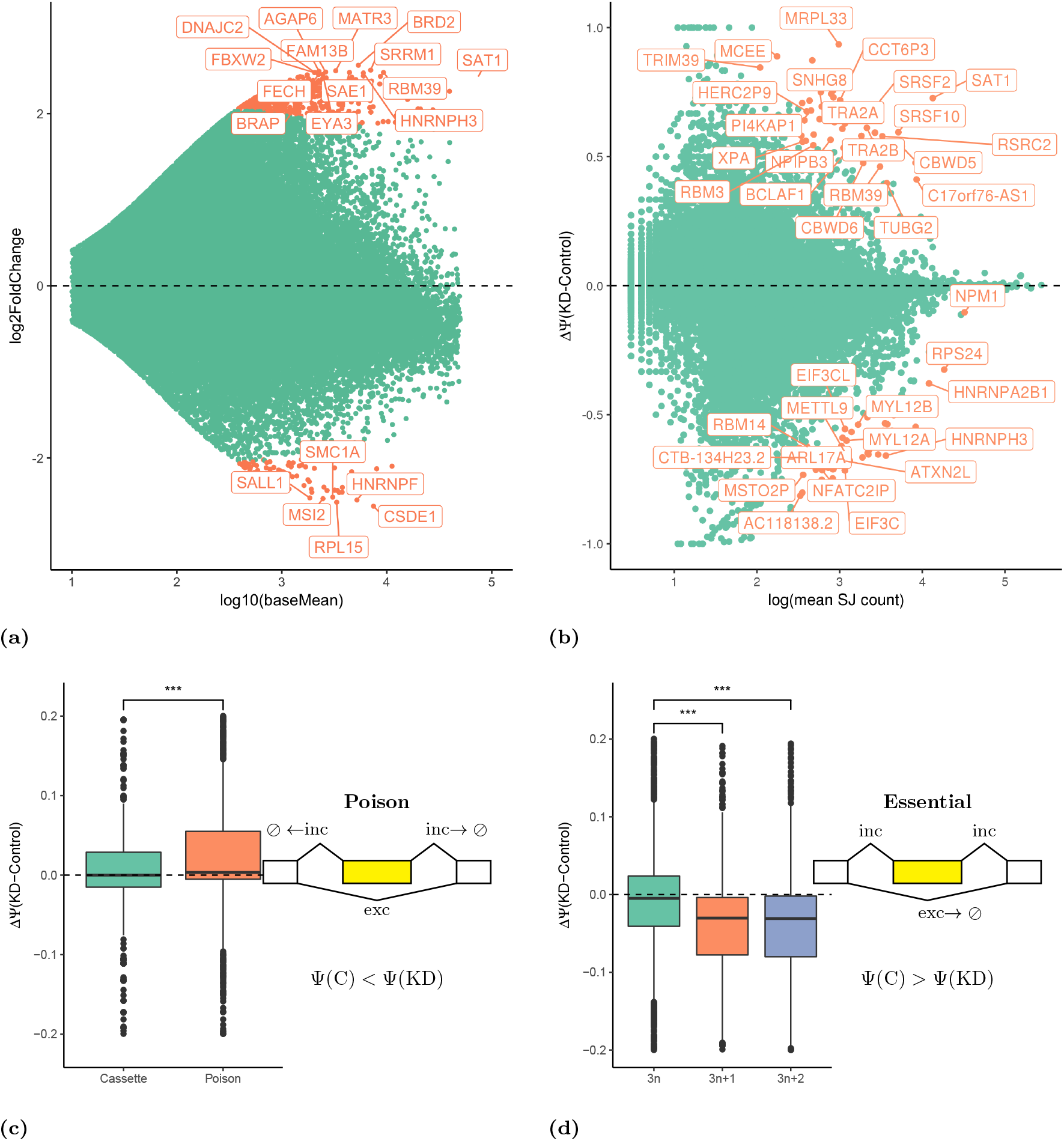
**(a)** DEseq2 analysis of differential gene expression in UPF1/XRN1 co-depletion. The top 15 overexpressed genes are shown in Supplementary Table S3. **(b)** Differential exon inclusion analysis in UPF1/XRN1 co-depletion. Changes at the 1% significance level are shown in orange. The list of differential exons is shown in Supplementary Data File 2. **(c)** Poison exons increase their inclusion rate upon UPF1/XRN1 co-depletion. **(d)** Essential exons (3*n* + 1 and 3*n* + 2) decrease their inclusion rate upon UPF1/XRN1 co-depletion, while non-essential (3*n*) exons remain on average unchanged.

In contrast, the analysis at the exon level (Figure 2b) reveals that a balanced proportion of exons significantly increase and decrease their level of inclusion when UPF1 and XRN1 are depleted (such exons are referred to as reactive exons). Among genes containing exons that respond to UPF1/XRN1 co-depletion again there are core spliceosomal proteins, serine-arginine rich proteins, hnRNPs, and general RNA-binding proteins (see Supplementary Table S5 for Gene Ontology analysis, and Supplementary Data File 2 for the list of ΔΨ values of reactive exons). Reactive exons are not uniformly distributed along transcripts: the 5’-exons and the 3’-exons are almost fivefold more likely to be reactive than are internal exons (Supplementary Figure S2). This result is consistent with the observation in yeast that NMD frequently affects splicing events in uORFs and 3’-UTRs [11].

### Poison and essential exons react oppositely to UPF1/XRN1 knockdown

In a broad sense, poison (or essential) exons are defined as exons that trigger NMD when included in the mRNA (or skipped, respectively). We asked whether poison and essential exons react differently to UPF1/XRN1 co-depletion. When NMD pathway is not perturbed, transcripts that contain poison exons are NMD targets. Hence, Ψ value of a poison exon should increase upon UPF1/XRN1 co-depletion because splice junctions (SJ) that support exon inclusion are degraded to a lesser extent when NMD is suppressed (Figure 2c). As expected, the distribution of ΔΨ for poison exons is significantly biased towards positive values as compared to the distribution of ΔΨ for the control set of non-poisonous cassette exons. For example, the inclusion of a poison exon that is located in the 3’-UTR of SRSF3 gene (ENST00000477442) is significantly upregulated in UPF1/XRN1 knockdown with ΔΨ = 0.39. Note that not all annotated poison exons react positively to UPF1/XRN1 co-depletion, likely reflecting complex higher-order responses in the gene regulatory network upon NMD pathway perturbation.

Conversely, SJs supporting the exclusion of an essential exon correspond to the transcripts that are degraded by NMD. Thus, the Ψ value of an essential exon should decrease upon UPF1/XRN1 co-depletion since exon skipping products are degraded to a lesser extent when NMD is suppressed (Figure 2d). The analysis of annotated exons shows that exons of length 3*n*, where *n* is an integer, generally do not introduce PTC, while exons of length 3*n* +1 and 3*n* + 2 lead to a frameshift that almost certainly induces a PTC (Supplementary Figure S1). That is, the majority of 3*n* +1 and 3*n* + 2 exons are essential, while the majority of 3*n* exons are non-essential. In accordance with this, the distribution of ΔΨ for 3*n* exons is symmetric around zero, while that of ΔΨ for 3*n* + 1 and 3*n* + 2 exons significantly deviates from zero towards negative values (Figure 2d). For example, skipping of exon 10 in PTBP2 is substantially downregulated upon UPF1/XRN1 co-depletion with ΔΨ = −0.67. Of note, the corresponding transcript isoform ENST00000541987 is not yet annotated as NMD.

### Poison and essential exons react oppositely to RBP perturbation

In this section, we focus on differential splicing analysis in shRNA-KD of a large panel of RPBs followed by RNA-seq provided by ENCODE consortium [51]. Perturbations of RBP expression must affect autoregulatory mechanisms that sense cellular RBP concentrations. To quantify splicing changes, we again used ΔΨ between KD and control experiments [51], assuming that exon inclusion changes are independent of gene expression changes, and that both mRNA and protein levels of RBP decrease with respect to their baseline levels after shRNA-KD.

Since RBP binding may exert both activating and repressing effects on exon inclusion, the autoregulation via AS-NMD can be achieved by either activating or repressing mechanisms. In the former case, the excess of RBP may activate the inclusion of a poison exon in its pre-mRNA, which would lead to the degradation of that pre-mRNA by NMD (Figure 3a). Then, shRNA-KD of an activating RBP should lead to a lower inclusion rate of the poison exon, i.e., ΔΨ of such exon should be negative. Conversely, the excess of RBP may suppress the inclusion of an essential exon, also leading to the degradation by NMD (Figure 3b). Then, shRNA-KD of a repressive RBP should result in a higher inclusion level of the essential exon, i.e., ΔΨ of such exon should be positive. Thus, poison and essential exons react oppositely to the perturbations of their host genes and to NMD pathway perturbations. These anticorrelated changes are illustrated in Figure 3c.

**Figure 3.**
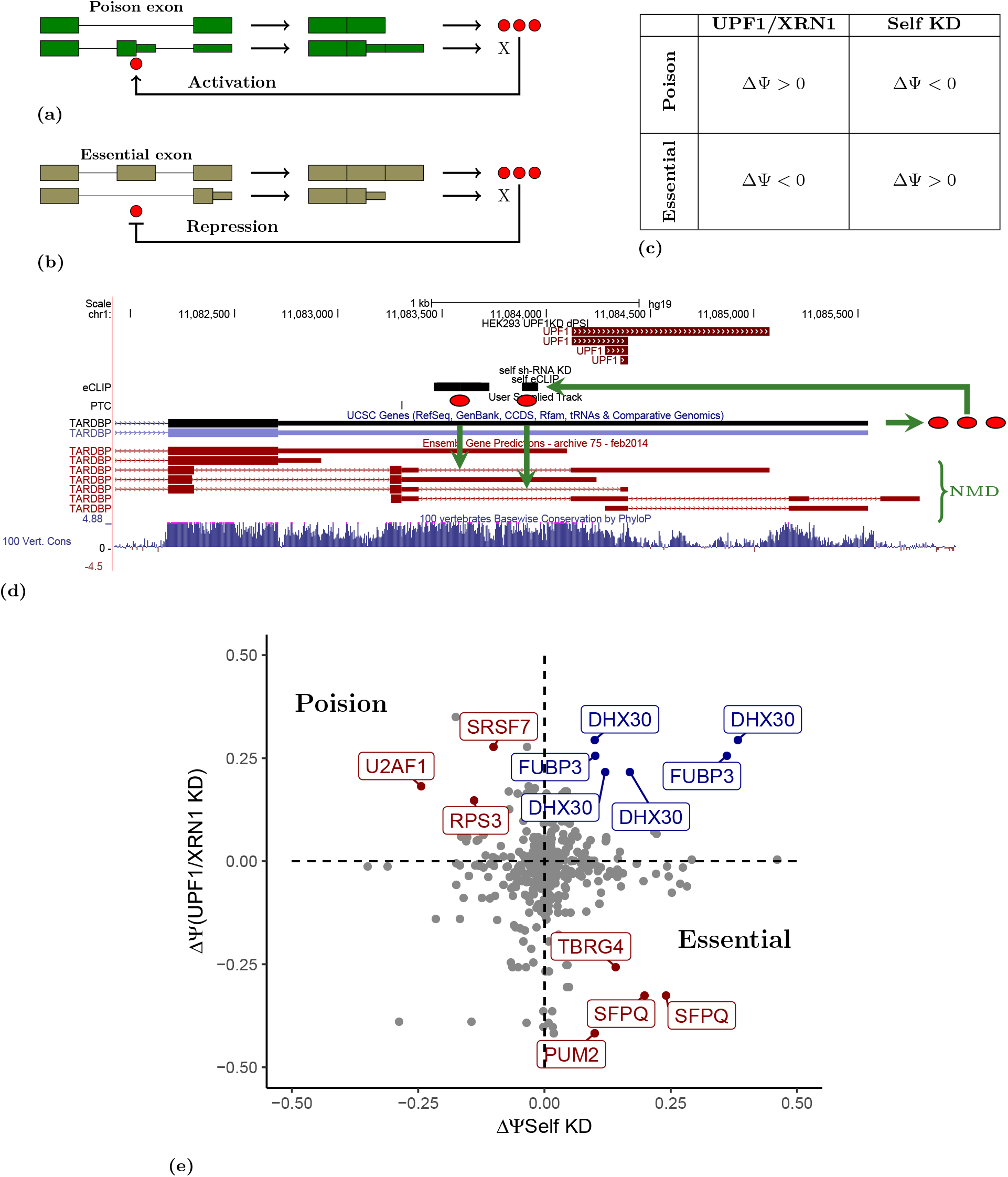
**(a)** RBP binding to its own pre-mRNA promotes inclusion of a poison exon and creates a negative feedback loop; shRNA-KD of an activating RBP should promote exon skipping. **(b)** RBP binding to its own pre-mRNA promotes skipping of an essential exon; shRNA-KD of a repressing RBP should promote exon inclusion. **(c)** A summary of the expected splicing changes in poison and essential exons under UPF1/XRN1 co-depletion and the depletion of RBP itself. **(d)** TARDBP protein binds its 3’-UTR (binding sites are shown in black in the eCLIP track) and promotes splicing at poison exons. **(e)** A scatterplot of significant splicing changes under UPF1/XRN1 co-depletion (y axis) vs. shRNA-KD of RBP itself (x-axis). Only genes with |ΔΨ| > 0.1 in both axes and with at least one cognate eCLIP peak are labelled. The labels are red for ΔΨ patterns from panel (c); the same-sign changes are shown by blue labels.

In order to apply this logic to the identification of novel AS-NMD autoregulatory targets, we first tested a few genes with known unproductive splicing. Studies have demonstrated that alternative skipping of exon 11 in Polypyrimidine Tract Binding Protein 1 (PTBP1) leads to an mRNA that is removed by NMD, and that this mechanism degrades a large part of the PTBP1 transcripts in HeLa cells [36]. While exon 11 is essential, the changes in its inclusion rate that are observed upon UPF1/XRN1 and PTBP1 depletion are small by the absolute value (ΔΨ_UPF1_ = –0.07 and ΔΨ_PTBP1_ = 0.025, respectively), but their opposite signs are consistent with the principle illustrated in Figure 3c. Interestingly, there is an eCLIP peak near exon 11 of PTBP1, but the two variants of exon 9 are reactive to PTBP1 depletion with much higher ΔΨ (Supplementary Figure S3) suggesting that skipping of exon 11 may be masked by nearby alternative splicing events.

Human TAR DNA-binding protein (TARDBP, TDP-43) is a key molecular player in amyotrophic lateral sclerosis and frontotemporal lobar degeneration. TARDBP controls its own expression level through a negative feedback loop, in which it binds to the 3’-UTR in its own mRNA and induces AS-NMD [31, 32]. Indeed, its 3’-UTR contains a number of unproductive splicing events that become activated upon UPF1/XRN1 co-depletion (Figure 3d, UPF1 track labelled in red, ΔΨ > 0). The 3’-UTR of TARDBP contains several cognate eCLIP peaks, which confirm that TARDBP is capable of binding its own 3’-UTR. However, there are no significant splicing changes in the 3’-UTR in response to TARDBP shRNA-KD (the expected change is ΔΨ < 0, see Figure 3c), which could be related to incomplete suppression of TARDBP by shRNA-KD. Indeed, according to gene expression quantification, the efficacy of shRNA suppression varies greatly between RBPs.

In order to obtain a list of high-confidence targets, we applied stringent thresholds to detect splicing changes that are not only significant, but also substantial by absolute value. Based on the published values [68], we used the cutoff of |ΔΨ| > 0.1 for both UPF1/XRN1 co-depletion and shRNA-KD of the RBP. Additionally, we imposed a requirement that the mRNA of the RBP must contain at least one eCLIP peak of the RBP itself. As a result, we obtained three candidate poison exons (SRSF7, U2AF1, and RPS3 genes), four candidate essential exons (in SFPQ, TBRG4, and PUM2 genes), and six exons that were upregulated in both UPF1/XRN1 co-depletion and shRNA-KD of the RBP. They are shown in Figure 3e by labelled points; genes with |ΔΨ| > 0.1 but with no eCLIP peaks are shown without labels.

In what follows, we use Genome Browser diagrams with custom tracks similar to ones shown in Figure 3d. The track labelled “UPF1” depicts ΔΨ, the exon inclusion change upon UPF1/XRN1 co-depletion, such that red color corresponds to ΔΨ *>* 0, and blue color corresponds to ΔΨ *<* 0. The track labelled with the name of the RBP shows ΔΨ for the shRNA-KD of the RBP itself using the same color code for ΔΨ. The black boxes in the track labelled “eCLIP” represent eCLIP peaks. For reader’s convenience, we use green arrows to indicate the sequence of events which, as we hypothesize, lead to AS-NMD. In the next section, we discuss potential AS-NMD autoregulatory mechanisms for these genes in detail.

## Case studies

### SRSF7

SRSF7, a classic member of serine/arginine-rich splicing factor family, is an important regulator of pre-mRNA splicing, nuclear export, and translation [69]. It has been reported recently that SRSF7 plays a major role in proliferation of cancer cells and apoptosis [70, 71]. Other serine/arginine-rich splicing factors, for example SRSF3, were shown to modulate their own alternative splicing, as well as that of other transcripts encoding SR proteins [72]. Here, we report a strong evidence for two poison exons in SRSF7 to be implicated in a negative autoregulatory feedback loop (Figure 4a). Indeed, exons 4a and 4b become substantially more included upon UPF1/XRN1 co-depletion (ΔΨ = 0.278 and ΔΨ = 0.165, respectively), confirming that they both are poison exons. On the other hand, the depletion of SRSF7 by shRNA-KD promotes skipping of these exons (ΔΨ = –0.10 for exon 4a and ΔΨ = –0.05 for exon 4b), which indicates that their inclusion was activated by SRSF7 itself. Additionally, there are two eCLIP peaks located in the two poison exons. Therefore, it appears plausible that the excess of SRSF7 protein binds its own pre-mRNA to promote exon 4a and exon 4b inclusion, thus regulating its level of expression via AS-NMD.

**Figure 4.**
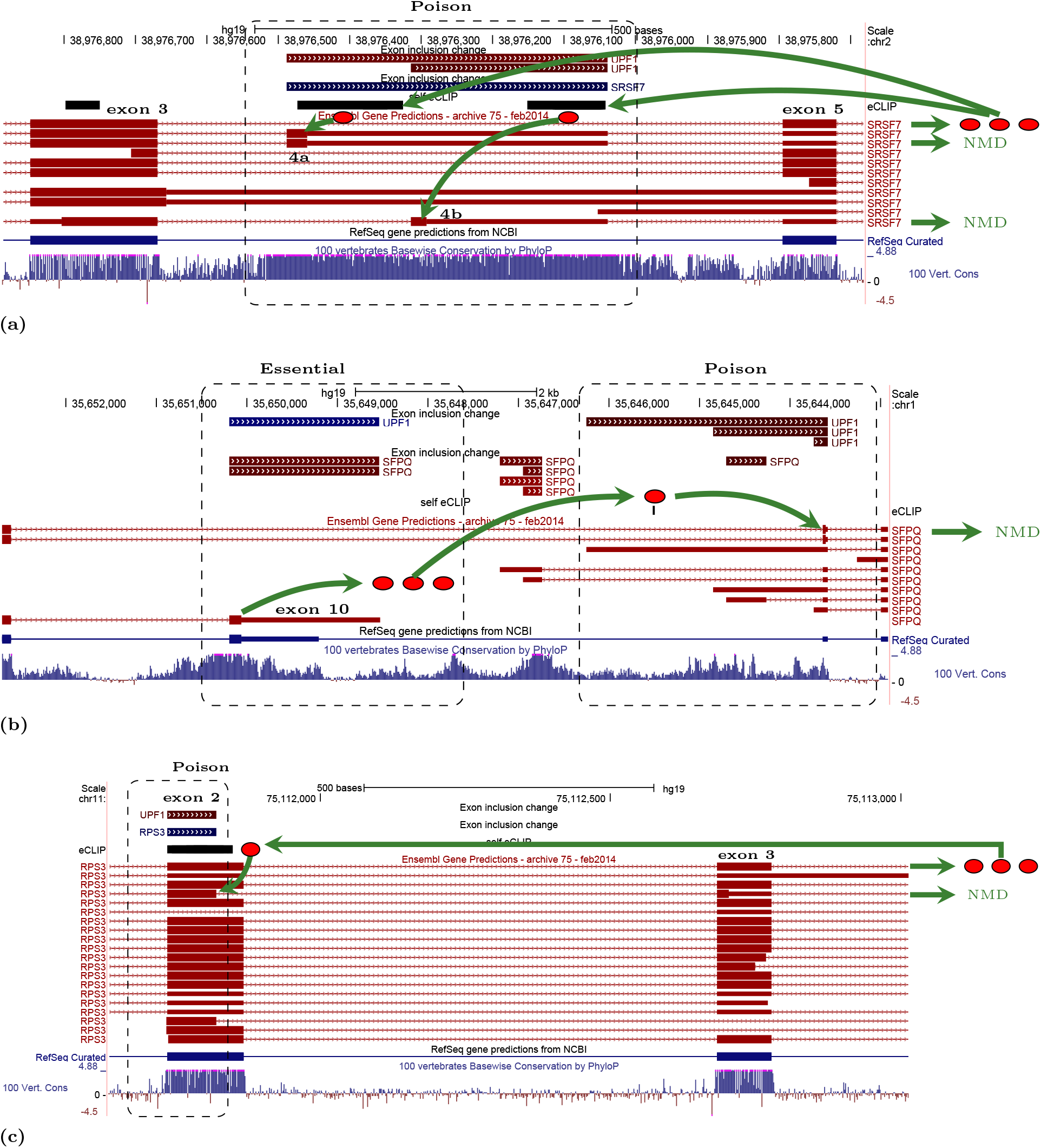
Case studies of poison and essential exons. The UPF1 track shows ΔΨ in UPF1/XRN1 co-depletion (ΔΨ *>* 0 are red, ΔΨ *<* 0 are blue). The track with the name of the RBP shows ΔΨ in RBP self-KD. The eCLIP track is shown in black. **(a)** SRSF7 binds its pre-mRNA to promote the inclusion of its poison exons 4a and 4b. **(b)** SFPQ binds its pre-mRNA co-transcriptionally and switches splicing towards the alternative 3’-UTR, which is an NMD substrate. **(c)** RPS3 binds its pre-mRNA around its exon 2 and suppresses splicing at the endogenous donor site by activating an upstream donor site, which leads to a frame shift and degradation by NMD.

### SFPQ

A member of another family of splicing factors, proline/glutamine-rich splicing factor SFPQ, is associated with amyotrophic lateral sclerosis [73], Alzheimer’s Disease and Frontotemporal Dementia [74]. We propose the following mechanism of SFPQ autoregulation by AS-NMD (Figure 4b). Exon 10, the terminal exon of the major SFPQ transcript isoform, is substantially downregulated in UPF1/XRN1 co-depletion (ΔΨ = –0.325), and it is also greatly upregulated when SFPQ itself is depleted (ΔΨ = 0.241), following the anticorrelated splicing pattern for essential exons (Figure 3c). We also noted that when exon 10 is suppressed, SFPQ splicing switches to a group of exons in the 3’-UTR, which are substantially upregulated in UPF1/XRN1 co-depletion, thus likely being poison exons. The intron spanning between exon 9 and the downstream poison exons contains a cognate eCLIP peak of SFPQ, indicating that alternative splicing and polyadenylation may be regulated by SFPQ itself. Indeed, examples of coupling between splicing and polyadenylation have been reported [75]. We therefore hypothesize that SFPQ binds its own pre-mRNA downstream of exon 10 and promotes alternative splicing to the distal 3’-UTR, which contains a PTC upstream of a splice junction and thus is a substrate of NMD.

### RPS3

Human ribosomal protein S3 (RPS3) is a component of the 40S ribosomal subunit that is mainly associated with protein synthesis. However, RPS3 has many additional extraribosomal functions and is involved in apoptosis and tumorigenesis [76]. Studies in *Caenorhabditis elegans* have shown that it is not uncommon among ribosomal proteins to be AS-NMD targets [25]. In particular, ribosomal proteins L3, L10a, and L12 use an evolutionarily-conserved pathway to insert PTCs in their pre-mRNA (76–78). Here we show that exon 2 of RPS3 has an alternative upstream donor site that induces a frameshift and targets RPS3 pre-mRNA for degradation (Figure 4c). This shorter variant of exon 2 reacts positively to UPF1/XRN1 co-depletion, confirming that the transcript with a frameshift is indeed a NMD target, and reacts negatively to RPS3 depletion, indicating that RPS3 is involved in promoting its inclusion. Consistently with this, exon 2 of RPS3 pre-mRNA contains a cognate eCLIP peak of RPS3, suggesting autoregulatory negative feedback loop with a poison 5’-splice site in this gene. Remarkably, the ribosomal protein L10a in *Caenorhabditis elegans* uses exactly the same strategy of regulation by specifically switching the donor site of its intron 3 to create an unproductively spliced mRNA [78].

### U2AF1

U2 small nuclear RNA auxiliary factor 1 (U2AF1) is another member of the serine/arginine-rich splicing factor family [79]. It encodes the small subunit of U2 auxiliary factor, a basic component of the major spliceosome which mediates binding of U2 snRNP to the pre-mRNA branch site (79–81). U2AF1 is frequently mutated in cancers, particularly in myelodysplastic syndromes, along with other mutated splicing factors [82]. In humans, exon 3 of U2AF1 exists in two mutually exclusive variants, exon 3a and exon 3b (Figure 5a). These exons are homologous (68.2% sequence identity) and have the same length of 67 nt, which suggests that they have evolved through a tandem genomic duplication [83]. Since 67 is not a multiple of three, the simultaneous inclusion of exons 3a and 3b, or simultaneous skipping of both, leads to a frameshift. Exon 3b, which is located downstream of exon 3a, reacts positively to UPF1/XRN1 co-depletion (ΔΨ = 0.18), and negatively to U2AF1 depletion (ΔΨ = –0.244), suggesting that it is, in fact, a poison exon. Indeed, the transcript isoform ENST00000464750 shows that exon 3b contains a PTC when included in the transcript together with exon 3a. Consistently with this, two eCLIP peaks are present downstream of exons 3a and 3b suggesting that U2AF1 binding may promote inclusion of both these exons, thereby creating a negative feedback loop of AS-NMD.

**Figure 5.**
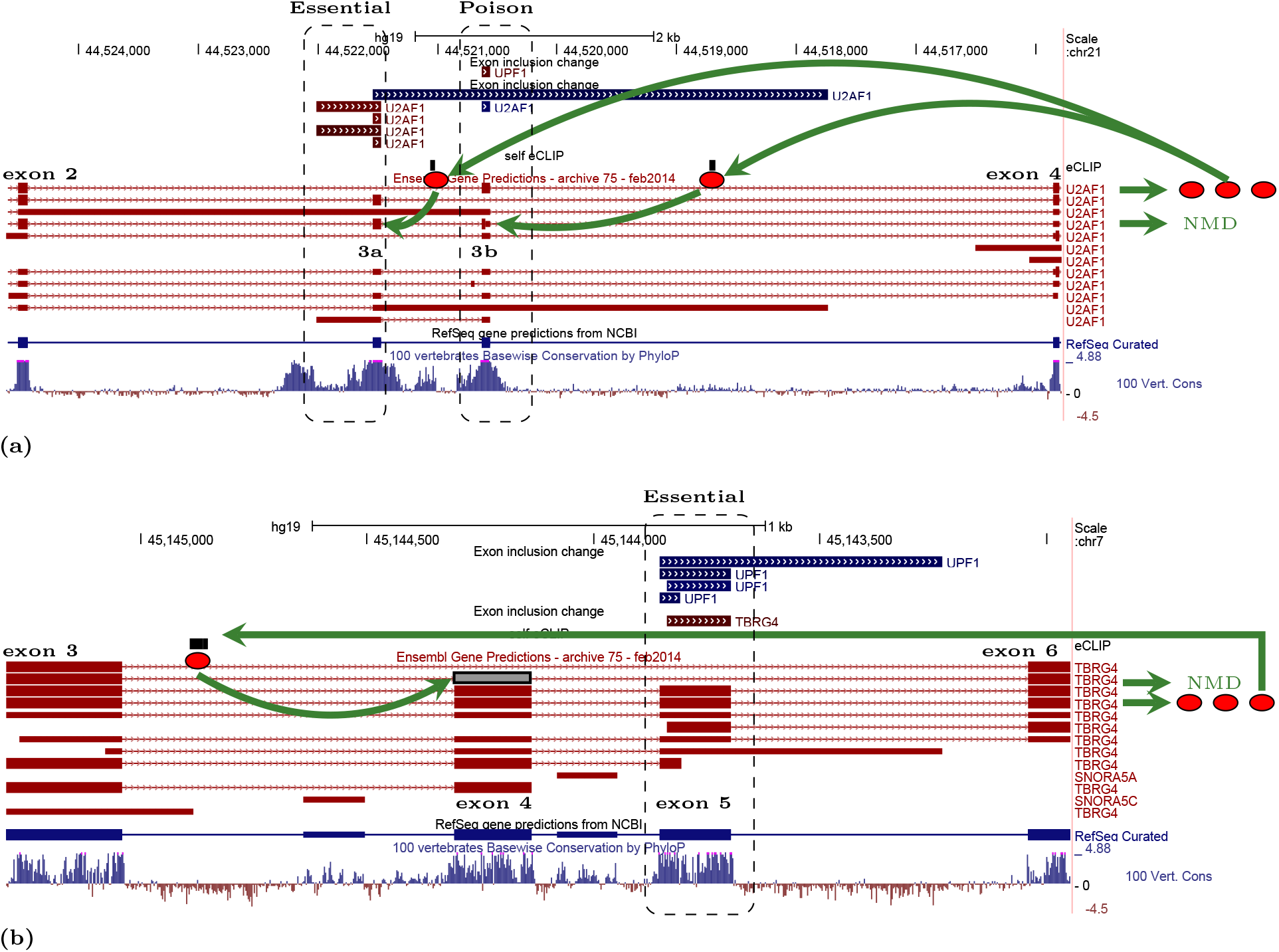
**(a)** U2AF1 contains two mutually exclusive variants of exon 3. If exon 3a is included, the inclusion of exon 3b leads to a frame shift and NMD. According to eCLIP, U2AF1 binds its pre-mRNA around exons 3a and 3b, which leads to their simultaneous inclusion, frame shift, and NMD. **(b)** Exons 4a and 4b of TBRG4 are mutually inclusive, i.e., either they are both included, or both skipped. Inclusion of each of them alone leads to a frame shift and NMD. eCLIP suggests that TBRG4 could bind its pre-mRNA upstream of exon 3a to facilitate its inclusion, thereby leading to NMD.

### TBRG4

Transforming growth factor beta regulator 4 (TBRG4), previously called FAST kinase domain-containing protein 4 (FASTKD4), is a member of FASTK family of proteins which are involved in the regulation of energy balance and RNA homeostasis in mitochondria [84]. TBRG4 is associated with cell proliferation in hematopoietic lineage and lung cancers [85]. Exons 4 and 5 of human TBRG4 are 172-nt and 158-nt long, respectively, and have a moderate sequence identity (Figure 5b). Unlike the case of U2AF1, exons 4 and 5 of TBRG4 are mutually inclusive, i.e., either they are both included, or both skipped, because the inclusion of each of them alone leads to a frameshift and NMD. Exon 5 responds negatively to UPF1/XRN1 co-depletion (ΔΨ = –0.256) and becomes more included when TBRG4 is depleted (ΔΨ = 0.141). According to eCLIP, TBRG4 binds its own pre-mRNA upstream of exon 4. We hypothesize that TBRG4 binding promotes inclusion of exon 4 and skipping of exon 5 to downregulate its own expression level by AS-NMD.

## Other genes

The most upstream exon in the beginning of the 5’-UTR of PUM2, human Pumilio homolog 2 gene, is likely to be essential, since its inclusion is suppressed in UPF1/XRN1 co-depletion and activated when PUM2 itself is downregulated (Figure 3e). According to eCLIP, shortly downstream of this exon there is a PUM2 binding site. This data suggest that the expression of this gene may be affected by an uORF, and the inclusion of the most upstream exon is needed to suppress the translation of this uORF (Supplementary Figure S4). Of note, the 3’-UTR of PUM2 contains a large cluster of its cognate binding sites, indicating that this gene has an additional AS-NMD control region.

Among our candidates there are exons that react positively to both UPF1/XRN1 co-depletion and to that of the host gene (Figure 3e). This pattern corresponds to a positive feedback loop (suppression of a poison exon or activation of an essential exon) rather than to the opposite ΔΨ changes characteristic for a negative feedback loop. In particular, these are exons 5 and 6 of the RNA helicase DHX30, a member of DEAD box family of proteins which are characterized by the conserved motif Asp-Glu-Ala-Asp (DEAD), and exons 13b and 17 of FUBP3, the Far Upstream Element Binding Protein 3. However, in both these cases the eCLIP track contains no evidence of cognate binding sites close to the regulatory exons. We therefore did not pursue the analysis of DHX30 and FUBP3 any further.

## Discussion

Gene expression includes a wide range of regulatory mechanisms that are used by the cells to adjust the production of a specific gene in response to various inputs [1]. All steps of gene expression, from the transcription initiation to the post-translational protein modification, are modulated within a sophisticated gene regulatory network, in which one regulator controls, and is itself controlled, by the expression of multiple other genes. A frequently observed pattern in such networks is the so-called self-loop, i.e., an autoregulatory feedback of a gene onto itself. Autoregulation provides a simple and, perhaps, the most robust regulatory feedback that doesn’t require any intermediate steps, allowing to sense directly the cellular concentration of a given factor. More than a half of the transcription factors in bacteria regulate their own genes by a self-loop, in which a factor binds to its own promoter and either activates or represses the transcription [86, 87].

While some eukaryotic genes use autoregulation self-loops at the transcriptional level, the expression can also be modulated post-transcriptionally [88]. As an example, the binding of YBX1 to a regulatory element in the 3’-UTR of YBX1 mRNA selectively inhibits its own translation [89]. However, the major way to modulate gene expression post-transcriptionally is through affecting mRNA stability [90]. Particularly, AS-NMD is a mechanism of regulation by mRNA degradation, which generates transcript isoforms with PTCs and promotes mRNA elimination by NMD [90]. In order for it to work through a self-loop, the gene product should be able to bind its own pre-mRNA. It is therefore not completely unexpected that RPBs are enriched among GWN, as well as among genes that autoregulate their expression via unproductive splicing.

In this work, we presented for the first time a bioinformatic analysis of a large panel of high-throughput sequencing data with the aim to identify novel cases of AS-NMD self-loops. By using a conservative approach and choosing stringent thresholds (|ΔΨ| > 0.1 for shRNA-KD, and *logFC* > 3 and *P* < 10^−3^ for eCLIP), we identified a set of candidate exons that could be involved in the autoregulation of RBP expression. However, exons in genes with known autoregulatory negative feedback loops, such as PTBP1 and TARDBP, showed small ΔΨ changes compared to these cutoffs, although the direction of the changes was consistent with the expected behavior (Figure 3c). The major factor contributing to this discrepancy in ΔΨ is the efficacy of shRNA-KD, which varies greatly between RBPs, and in the worst case of inefficient shRNA-KD, we underestimate the magnitude of ΔΨ. We therefore expect that our approach should suffer more from the false negative rate than from the false positive rate, and that high ΔΨ values in the examples shown in Figures 4 and 5 provide sufficient evidence to be high-confidence candidates for experimental validation. Regarding false positive predictions, a possibility remains that the observed ΔΨ, even as large by absolute value, could result from indirect responses in gene regulatory networks; however, we expect that these confounding effects are minimized by using several independent data sources.

Besides the efficacy of the gene knockdown, a number of other factors confound our analysis, including the specificity and efficacy of cross-linking and immunoprecipitation (IP). The efficacy of RBP-RNA interaction assessment depends on the crosslinking method and varies for single-stranded and double-stranded RNAs [91]. Besides this, the crosslinking position could itself be confounded by intramolecular base pairings, which often play a role in RNA processing [92], or some proteins within the size-matched control fraction of eCLIP could be not completely purified away [91]. Also, it cannot be excluded that IP sample is contaminated by cross-linked interacting RPBs because they often control their targets combinatorially through interacting with closely located binding sites, or that multiple domains of the same factor independently make contacts with distinct portions of the pre-mRNA and the actual binding site could be different from the crosslinking position [93]. As a result, eCLIP track may contain false positive peaks that represent the binding of the interacting partners or, conversely, some of the true eCLIP peaks could be missing.

According to Figure 3c, a negative feedback AS-NMD loop is characterized by the opposite reaction of its regulatory exons, poison or essential, to the inactivation of NMD pathway and to that of the RBP itself. Exons that react positively to both these perturbations suggest the existence of a similar positive feedback mechanism, which would work through repression of a poisonous exon, or activation of an essential exon in a manner opposite to that shown in Figures 3a and 3b. Unlike negative feedback loops, which tend to stabilize the output of a gene regulatory circuit by compensatory changes in the direction opposite to the original deviation, positive feedback systems are less common, but have other features that are important in biological systems, including bistability, hysteresis and non-linear activation properties [94]. An example of a positive feedback loop architecture at the transcriptional level is the regulatory network of four TFs of the bacterial DtxR family that maintains intracellular iron balance in archea [95]. Considering that the global architecture of SF regulatory networks is quite different from that of transcription factors [45], with more of a homeostatic role for SFs and more of a differentiating role for transcription factors, it remains an open question whether positive feedback loops at the post-transcriptional level exist at all. To date, there is no evidence of a positive feedback loop that works through AS-NMD, and the candidates studied here (DHX30 and FBP3) do not represent any convincing mechanism.

The approach implemented here is based on the analysis of exon inclusion rates in two particular cases of poison and essential exons. We used the definition of exon inclusion rate Ψ that was originally introduced for cassette exons [60], but can also be interpreted in a broader sense for a larger class of splicing events, including alternative donor and acceptor site usage. A similar analysis of the Completeness of Splicing Index [60] can be used to identify regulatory intron retention cases, which are often associated with down-regulation of gene expression via NMD [96]. In general, this methodology can be applied to arbitrary types of splicing events that are associated with downstream PTCs. However, such splicing events are often missing from the annotation databases because of undercoverage bias as a result of NMD degradation, and the computational identification of NMD transcripts is not completely straightforward. Similarly, the same approach could be applied to the discovery of cross-regulation of RBP expression, in conjunction with the analysis presented by Desai *et al* [45], but this and many other follow-up questions are beyond the scope of this short report.

## Conclusion

We presented a bioinformatic analysis of a large panel of high-throughput binding and gene expression assays in order to identify novel autoregulatory feedback loops of alternative splicing coupled with nonsense-mediated mRNA decay. By using stringent thresholds, we identified five candidate genes, SRSF7, SFPQ, RPS3, U2AF1, and TBRG4, in which the arrangement of binding sites and the responses to shRNA perturbations suggest clear mechanistic scenarios of gene regulation by NMD. It is likely that many other RBP use the same strategy of maintaining their physiological concentrations via alternative splicing and NMD. The framework outlined here can be used as a generic strategy for computational identification of post-transcriptional gene regulatory networks that operate through nonsense mediated mRNA decay.

## Supporting information

## Acknowledgements

We thank Zoya Chervontseva for critical comments and in-depth reading the manuscript. The work of DP and YP was supported by Skolkovo Institute of Science and Technology Research Grant RF-0000000653 and by Russian Foundation for Basic Research grant 18–29–13020-MK. AF and AB were supported by National Human Genome Research Institute of the National Institutes of Health under Award Number U41HG007234. The content is solely the responsibility of the authors and does not necessarily represent the official views of the National Institutes of Health. Work in the laboratory of RG was supported by the National Human Genome Research Institute of the US National Institutes of Health (grants U41HG007000, U54HG007004 and 1U24HG009446). BB was supported by a fellowship from la Secretaría de Universidades e Investigation del Departamento de Economía y Conocimiento de la Generalidad de Cataluña. We acknowledge support of the Spanish Ministry of Economy, Industry and Competitiveness (MEIC) to the EMBL partnership, to the Spanish Ministry of Economy and Competitiveness (MEC) ‘Centro de Excelencia Severo Ochoa’, and to the CERCA Programme/Generalitat de Catalunya.

## Author’s contributions

DP, AF, and RG conceived the analysis; DP designed the analysis; DP, YP, AB, and BB performed the analysis; all authors wrote the paper.

## References

1. Borbolis, F. and Syntichaki, P. (Dec, 2015) Cytoplasmic mRNA turnover and ageing. Mech. Ageing Dev., 152, 32–42.

2. Dassi, E. (2017) Handshakes and Fights: The Regulatory Interplay of RNA-Binding Proteins. Front Mol Biosci, 4, 67.

3. Baker, K. E. and Parker, R. (Jun, 2004) Nonsense-mediated mRNA decay: terminating erroneous gene expression. Curr. Opin. Cell Biol., 16(3), 293–299.

4. Karousis, E. D., Nasif, S., and Muhlemann, O. (09, 2016) Nonsense-mediated mRNA decay: novel mechanistic insights and biological impact. Wiley Interdiscip Rev RNA, 7(5), 661–682.

5. Popp, M. W. and Maquat, L. E. (Jun, 2016) Leveraging Rules of Nonsense-Mediated mRNA Decay for Genome Engineering and Personalized Medicine. Cell, 165(6), 1319–1322.

6. Fang, Y., Bateman, J. F., Mercer, J. F., and Lamande, S. R. (Jun, 2013) Nonsense-mediated mRNA decay of collagen-emerging complexity in RNA surveillance mechanisms. J. Cell. Sci., 126(Pt 12), 2551–2560.

7. Lewis, B. P., Green, R. E., and Brenner, S. E. (Jan, 2003) Evidence for the widespread coupling of alternative splicing and nonsense-mediated mRNA decay in humans. Proc. Natl. Acad. Sci. U.S.A., 100(1), 189–192.

8. Lareau, L. F., Brooks, A. N., Soergel, D. A., Meng, Q., and Brenner, S. E. (2007) The coupling of alternative splicing and nonsense-mediated mRNA decay. Adv. Exp. Med. Biol., 623, 190–211.

9. Tsyba, L., Skrypkina, I., Rynditch, A., Nikolaienko, O., Ferenets, G., Fortna, A., and Gardiner, K. (Jul, 2004) Alternative splicing of mammalian Intersectin 1: domain associations and tissue specificities. Genomics, 84(1), 106–113.

10. Pacheco, T. R., Gomes, A. Q., Barbosa-Morais, N. L., Benes, V., Ansorge, W., Wollerton, M., Smith, C. W., Valcarcel, J., and Carmo-Fonseca, M. (Jun, 2004) Diversity of vertebrate splicing factor U2AF35: identification of alternatively spliced U2AF1 mRNAS. J. Biol. Chem., 279(26), 27039–27049.

11. Kebaara, B. W. and Atkin, A. L. (May, 2009) Long 3’-UTRs target wild-type mRNAs for nonsense-mediated mRNA decay in Saccharomyces cerevisiae. Nucleic Acids Res., 37(9), 2771–2778.

12. Mendell, J. T., Sharifi, N. A., Meyers, J. L., Martinez-Murillo, F., and Dietz, H. C. (Oct, 2004) Nonsense surveillance regulates expression of diverse classes of mammalian transcripts and mutes genomic noise. Nat. Genet., 36(10), 1073–1078.

13. Ni, J. Z., Grate, L., Donohue, J. P., Preston, C., Nobida, N., O’Brien, G., Shiue, L., Clark, T. A., Blume, J. E., and Ares, M. (Mar, 2007) Ultraconserved elements are associated with homeostatic control of splicing regulators by alternative splicing and nonsense-mediated decay. Genes Dev., 21(6), 708–718.

14. Lareau, L. F., Inada, M., Green, R. E., Wengrod, J. C., and Brenner, S. E. (Apr, 2007) Unproductive splicing of SR genes associated with highly conserved and ultraconserved DNA elements. Nature, 446(7138), 926–929.

15. Lareau, L. F. and Brenner, S. E. (Apr, 2015) Regulation of splicing factors by alternative splicing and NMD is conserved between kingdoms yet evolutionarily flexible. Mol. Biol. Evol., 32(4), 1072–1079.

16. Kalyna, M., Simpson, C. G., Syed, N. H., Lewandowska, D., Marquez, Y., Kusenda, B., Marshall, J., Fuller, J., Cardle, L., McNicol, J., Dinh, H. Q., Barta, A., and Brown, J. W. (Mar, 2012) Alternative splicing and nonsense-mediated decay modulate expression of important regulatory genes in Arabidopsis. Nucleic Acids Res., 40(6), 2454–2469.

17. Mudge, J. M., Frankish, A., and Harrow, J. (Dec, 2013) Functional transcriptomics in the post-ENCODE era. Genome Res., 23(12), 1961–1973.

18. Popp, M. W. and Maquat, L. E. (Feb, 2018) Nonsense-mediated mRNA Decay and Cancer. Curr. Opin. Genet. Dev., 48, 44–50.

19. Munoz, U., Puche, J. E., Hannivoort, R., Lang, U. E., Cohen-Naftaly, M., and Friedman, S. L. (Sep, 2012) Hepatocyte growth factor enhances alternative splicing of the Kruppel-like factor 6 (KLF6) tumor suppressor to promote growth through SRSF1. Mol. Cancer Res., 10(9), 1216–1227.

20. Wang, D., Zavadil, J., Martin, L., Parisi, F., Friedman, E., Levy, D., Harding, H., Ron, D., and Gardner, L. B. (Sep, 2011) Inhibition of nonsense-mediated RNA decay by the tumor microenvironment promotes tumorigenesis. Mol. Cell. Biol., 31(17), 3670–3680.

21. Lykke-Andersen, S. and Jensen, T. H. (Nov, 2015) Nonsense-mediated mRNA decay: an intricate machinery that shapes transcriptomes. Nat. Rev. Mol. Cell Biol., 16(11), 665–677.

22. Smith, J. E., Alvarez-Dominguez, J. R., Kline, N., Huynh, N. J., Geisler, S., Hu, W., Coller, J., and Baker, K. E. (Jun, 2014) Translation of small open reading frames within unannotated RNA transcripts in Saccharomyces cerevisiae. Cell Rep, 7(6), 1858–1866.

23. Hurt, J. A., Robertson, A. D., and Burge, C. B. (Oct, 2013) Global analyses of UPF1 binding and function reveal expanded scope of nonsense-mediated mRNA decay. Genome Res., 23(10), 1636–1650.

24. Li, L., Geng, Y., Feng, R., Zhu, Q., Miao, B., Cao, J., and Fei, S. (2017) The Human RNA Surveillance Factor UPF1 Modulates Gastric Cancer Progression by Targeting Long Non-Coding RNA MALAT1. Cell. Physiol. Biochem., 42(6), 2194–2206.

25. Mitrovich, Q. M. and Anderson, P. (Sep, 2000) Unproductively spliced ribosomal protein mRNAs are natural targets of mRNA surveillance in C. elegans. Genes Dev., 14(17), 2173–2184.

26. Long, J. C. and Caceres, J. F. (Jan, 2009) The SR protein family of splicing factors: master regulators of gene expression. Biochem. J., 417(1), 15–27.

27. Sun, S., Zhang, Z., Sinha, R., Karni, R., and Krainer, A. R. (Mar, 2010) SF2/ASF autoregulation involves multiple layers of post-transcriptional and translational control. Nat. Struct. Mol. Biol., 17(3), 306–312.

28. Han, S. P., Tang, Y. H., and Smith, R. (Sep, 2010) Functional diversity of the hnRNPs: past, present and perspectives. Biochem. J., 430(3), 379–392.

29. McGlincy, N. J., Tan, L. Y., Paul, N., Zavolan, M., Lilley, K. S., and Smith, C. W. (Oct, 2010) Expression proteomics of UPF1 knockdown in HeLa cells reveals autoregulation of hnRNP A2/B1 mediated by alternative splicing resulting in nonsense-mediated mRNA decay. BMC Genomics, 11, 565.

30. Rossbach, O., Hung, L. H., Schreiner, S., Grishina, I., Heiner, M., Hui, J., and Bindereif, A. (Mar, 2009) Auto- and cross-regulation of the hnRNP L proteins by alternative splicing. Mol. Cell. Biol., 29(6), 1442–1451.

31. Polymenidou, M., Lagier-Tourenne, C., Hutt, K. R., Huelga, S. C., Moran, J., Liang, T. Y., Ling, S. C., Sun, E., Wancewicz, E., Mazur, C., Kordasiewicz, H., Sedaghat, Y., Donohue, J. P., Shiue, L., Bennett, C. F., Yeo, G. W., and Cleveland, D. W. (Apr, 2011) Long pre-mRNA depletion and RNA missplicing contribute to neuronal vulnerability from loss of TDP-43. Nat. Neurosci., 14(4), 459–468.

32. Ayala, Y. M., De Conti, L., Avendano-Vazquez, S. E., Dhir, A., Romano, M., D’Ambrogio, A., Tollervey, J., Ule, J., Baralle, M., Buratti, E., and Baralle, F. E. (Jan, 2011) TDP-43 regulates its mRNA levels through a negative feedback loop. EMBO J., 30(2), 277–288.

33. Stoilov, P., Daoud, R., Nayler, O., and Stamm, S. (Mar, 2004) Human tra2-beta1 autoregulates its protein concentration by influencing alternative splicing of its pre-mRNA. Hum. Mol. Genet., 13(5), 509–524.

34. Gates, D. P., Coonrod, L. A., and Berglund, J. A. (Sep, 2011) Autoregulated splicing of muscleblind like 1 (MBNL1) Pre-mRNA. J. Biol. Chem., 286(39), 34224–34233.

35. Konieczny, P., Stepniak-Konieczna, E., and Sobczak, K. (01, 2018) MBNL expression in autoregulatory feedback loops. RNA Biol, 15(1), 1–8.

36. Wollerton, M. C., Gooding, C., Wagner, E. J., Garcia-Blanco, M. A., and Smith, C. W. (Jan, 2004) Autoregulation of polypyrimidine tract binding protein by alternative splicing leading to nonsense-mediated decay. Mol. Cell, 13(1), 91–100.

37. Izumikawa, K., Yoshikawa, H., Ishikawa, H., Nobe, Y., Yamauchi, Y., Philipsen, S., Simpson, R. J., Isobe, T., and Takahashi, N. (Nov, 2016) Chtop (Chromatin target of Prmt1) auto-regulates its expression level via intron retention and nonsense-mediated decay of its own mRNA. Nucleic Acids Res., 44(20), 9847–9859.

38. Zhou, Y., Liu, S., Liu, G., Ozturk, A., and Hicks, G. G. (Oct, 2013) ALS-associated FUS mutations result in compromised FUS alternative splicing and autoregulation. PLoS Genet., 9(10), e1003895.

39. Sun, Y., Bao, Y., Han, W., Song, F., Shen, X., Zhao, J., Zuo, J., Saffen, D., Chen, W., Wang, Z., You, X., and Wang, Y. (Aug, 2017) Autoregulation of RBM10 and cross-regulation of RBM10/RBM5 via alternative splicing-coupled nonsense-mediated decay. Nucleic Acids Res., 45(14), 8524–8540.

40. Saltzman, A. L., Kim, Y. K., Pan, Q., Fagnani, M. M., Maquat, L. E., and Blencowe, B. J. (Jul, 2008) Regulation of multiple core spliceosomal proteins by alternative splicing-coupled nonsense-mediated mRNA decay. Mol. Cell. Biol., 28(13), 4320–4330.

41. Cuccurese, M., Russo, G., Russo, A., and Pietropaolo, C. (2005) Alternative splicing and nonsense-mediated mRNA decay regulate mammalian ribosomal gene expression. Nucleic Acids Res., 33(18), 5965–5977.

42. Spellman, R., Llorian, M., and Smith, C. W. (Aug, 2007) Crossregulation and functional redundancy between the splicing regulator PTB and its paralogs nPTB and ROD1. Mol. Cell, 27(3), 420–434.

43. Jangi, M., Boutz, P. L., Paul, P., and Sharp, P. A. (Mar, 2014) Rbfox2 controls autoregulation in RNA-binding protein networks. Genes Dev., 28(6), 637–651.

44. Dredge, B. K. and Jensen, K. B. (2011) NeuN/Rbfox3 nuclear and cytoplasmic isoforms differentially regulate alternative splicing and nonsense-mediated decay of Rbfox2. PLoS ONE, 6(6), e21585.

45. Desai, A., Hu, Z., French, C. E., Lloyd, J. P. B., and Brenner, S. E. (2018) Networks of Splice Factor Regulation by Unproductive Splicing Coupled With NMD. Genome Biology, submitted.

46. Kino, Y., Washizu, C., Kurosawa, M., Oma, Y., Hattori, N., Ishiura, S., and Nukina, N. (Feb, 2015) Nuclear localization of MBNL1: splicing-mediated autoregulation and repression of repeat-derived aberrant proteins. Hum. Mol. Genet., 24(3), 740–756.

47. Budini, M. and Buratti, E. (Nov, 2011) TDP-43 autoregulation: implications for disease. J. Mol. Neurosci., 45(3), 473–479.

48. Guo, J., Jia, J., and Jia, R. (Sep, 2015) PTBP1 and PTBP2 impaired autoregulation of SRSF3 in cancer cells. Sci Rep, 5, 14548.

49. Jacob, A. G. and Smith, C. W. (09, 2017) Intron retention as a component of regulated gene expression programs. Hum. Genet., 136(9), 1043–1057.

50. Van Nostrand, E. L., Pratt, G. A., Shishkin, A. A., Gelboin-Burkhart, C., Fang, M. Y., Sundararaman, B., Blue, S. M., Nguyen, T. B., Surka, C., Elkins, K., Stanton, R., Rigo, F., Guttman, M., and Yeo, G. W. (06, 2016) Robust transcriptome-wide discovery of RNA-binding protein binding sites with enhanced CLIP (eCLIP). Nat. Methods, 13(6), 508–514.

51. Van Nostrand, E. L., Freese, P., Pratt, G. A., Wang, X., Wei, X., Blue, S. M., Dominguez, D., Cody, N. A. L., Olson, S., Sundararaman, B., Xiao, R., Zhan, L., Bazile, C., Benoit Bouvrette, L. P., Chen, J., Duff, M. O., Garcia, K., Gelboin-Burkhart, C., Hochman, A., Lambert, N. J., Li, H., Nguyen, T. B., Palden, T., Rabano, I., Sathe, S., Stanton, R., Louie, A. L., Aigner, S., Bergalet, J., Zhou, B., Su, A., Wang, R., Yee, B. A., Fu, X.-D., Lecuyer, E., Burge, C. B., Graveley, B., and Yeo, G. W. (2017) A Large-Scale Binding and Functional Map of Human RNA Binding Proteins. bioRxiv,.

52. Consortium, T. E. (Sep, 2012) An integrated encyclopedia of DNA elements in the human genome. Nature, 489(7414), 57–74.

53. Sloan, C. A., Chan, E. T., Davidson, J. M., Malladi, V. S., Strattan, J. S., Hitz, B. C., Gabdank, I., Narayanan, A. K., Ho, M., Lee, B. T., Rowe, L. D., Dreszer, T. R., Roe, G., Podduturi, N. R., Tanaka, F., Hong, E. L., and Cherry, J. M. (Jan, 2016) ENCODE data at the ENCODE portal. Nucleic Acids Res., 44(D1), D726–732.

54. Lykke-Andersen, S., Chen, Y., Ardal, B. R., Lilje, B., Waage, J., Sandelin, A., and Jensen, T. H. (Nov, 2014) Human nonsense-mediated RNA decay initiates widely by endonucleolysis and targets snoRNA host genes. Genes Dev., 28(22), 2498–2517.

55. Church, D. M., Schneider, V. A., Graves, T., Auger, K., Cunningham, F., Bouk, N., Chen, H. C., Agarwala, R., McLaren, W. M., Ritchie, G. R., Albracht, D., Kremitzki, M., Rock, S., Kotkiewicz, H., Kremitzki, C., Wollam, A., Trani, L., Fulton, L., Fulton, R., Matthews, L., Whitehead, S., Chow, W., Torrance, J., Dunn, M., Harden, G., Threadgold, G., Wood, J., Collins, J., Heath, P., Griffiths, G., Pelan, S., Grafham, D., Eichler, E. E., Weinstock, G., Mardis, E. R., Wilson, R. K., Howe, K., Flicek, P., and Hubbard, T. (Jul, 2011) Modernizing reference genome assemblies. PLoS Biol., 9(7), e1001091.

56. Harrow, J., Frankish, A., Gonzalez, J. M., Tapanari, E., Diekhans, M., Kokocinski, F., Aken, B. L., Barrell, D., Zadissa, A., Searle, S., Barnes, I., Bignell, A., Boychenko, V., Hunt, T., Kay, M., Mukherjee, G., Rajan, J., Despacio-Reyes, G., Saunders, G., Steward, C., Harte, R., Lin, M., Howald, C., Tanzer, A., Derrien, T., Chrast, J., Walters, N., Balasubramanian, S., Pei, B., Tress, M., Rodriguez, J. M., Ezkurdia, I., van Baren, J., Brent, M., Haussler, D., Kellis, M., Valencia, A., Reymond, A., Gerstein, M., Guigo, R., and Hubbard, T. J. (Sep, 2012) GENCODE: the reference human genome annotation for The ENCODE Project. Genome Res., 22(9), 1760–1774.

57. Quinlan, A. R. and Hall, I. M. (Mar, 2010) BEDTools: a flexible suite of utilities for comparing genomic features. Bioinformatics, 26(6), 841–842.

58. authors listed, N. (01, 2017) Expansion of the Gene Ontology knowledgebase and resources. Nucleic Acids Res., 45(D1), D331–D338.

59. authors listed, N. (01, 2017) UniProt: the universal protein knowledgebase. Nucleic Acids Res., 45(D1), D158–D169.

60. Pervouchine, D. D., Knowles, D. G., and Guigo, R. (Jan, 2013) Intron-centric estimation of alternative splicing from RNA-seq data. Bioinformatics, 29(2), 273–274.

61. Trapnell, C., Roberts, A., Goff, L., Pertea, G., Kim, D., Kelley, D. R., Pimentel, H., Salzberg, S. L., Rinn, J. L., and Pachter, L. (Mar, 2012) Differential gene and transcript expression analysis of RNA-seq experiments with TopHat and Cufflinks. Nat Protoc, 7(3), 562–578.

62. Love, M. I., Huber, W., and Anders, S. (2014) Moderated estimation of fold change and dispersion for RNA-seq data with DESeq2. Genome Biol., 15(12), 550.

63. Ashburner, M., Ball, C. A., Blake, J. A., Botstein, D., Butler, H., Cherry, J. M., Davis, A. P., Dolinski, K., Dwight, S. S., Eppig, J. T., Harris, M. A., Hill, D. P., Issel-Tarver, L., Kasarskis, A., Lewis, S., Matese, J. C., Richardson, J. E., Ringwald, M., Rubin, G. M., and Sherlock, G. (May, 2000) Gene ontology: tool for the unification of biology. The Gene Ontology Consortium. Nat. Genet., 25(1), 25–29.

64. Falcon, S. and Gentleman, R. (Jan, 2007) Using GOstats to test gene lists for GO term association. Bioinformatics, 23(2), 257–258.

65. Eden, E., Navon, R., Steinfeld, I., Lipson, D., and Yakhini, Z. (Feb, 2009) GOrilla: a tool for discovery and visualization of enriched GO terms in ranked gene lists. BMC Bioinformatics, 10, 48.

66. Gatfield, D. and Izaurralde, E. (Jun, 2004) Nonsense-mediated messenger RNA decay is initiated by endonucleolytic cleavage in Drosophila. Nature, 429(6991), 575–578.

67. Franks, T. M., Singh, G., and Lykke-Andersen, J. (Dec, 2010) Upf1 ATPase-dependent mRNP disassembly is required for completion of nonsense-mediated mRNA decay. Cell, 143(6), 938–950.

68. McManus, C. J., Coolon, J. D., Eipper-Mains, J., Wittkopp, P. J., and Graveley, B. R. (May, 2014) Evolution of splicing regulatory networks in Drosophila. Genome Res., 24(5), 786–796.

69. Bourgeois, C. F., Lejeune, F., and Stevenin, J. (2004) Broad specificity of SR (serine/arginine) proteins in the regulation of alternative splicing of pre-messenger RNA. Prog. Nucleic Acid Res. Mol. Biol., 78, 37–88.

70. Fu, Y. and Wang, Y. (Apr, 2018) SRSF7 knockdown promotes apoptosis of colon and lung cancer cells. Oncol Lett, 15(4), 5545–5552.

71. Saijo, S., Kuwano, Y., Masuda, K., Nishikawa, T., Rokutan, K., and Nishida, K. (2016) Serine/arginine-rich splicing factor 7 regulates p21-dependent growth arrest in colon cancer cells. J. Med. Invest., 63(3–4), 219–226.

72. Anko, M. L., Muller-McNicoll, M., Brandl, H., Curk, T., Gorup, C., Henry, I., Ule, J., and Neugebauer, K. M. (2012) The RNA-binding landscapes of two SR proteins reveal unique functions and binding to diverse RNA classes. Genome Biol., 13(3), R17.

73. Luisier, R., Tyzack, G. E., Hall, C. E., Mitchell, J. S., Devine, H., Taha, D. M., Malik, B., Meyer, I., Greensmith, L., Newcombe, J., Ule, J., Luscombe, N. M., and Patani, R. (May, 2018) Intron retention and nuclear loss of SFPQ are molecular hallmarks of ALS. Nat Commun, 9(1), 2010.

74. Lu, J., Shu, R., and Zhu, Y. (2018) Dysregulation and Dislocation of SFPQ Disturbed DNA Organization in Alzheimer’s Disease and Frontotemporal Dementia. J. Alzheimers Dis., 61(4), 1311–1321.

75. Raker, V. A., Mironov, A. A., Gelfand, M. S., and Pervouchine, D. D. (Aug, 2009) Modulation of alternative splicing by long-range RNA structures in Drosophila. Nucleic Acids Res., 37(14), 4533–4544.

76. Han, S. H., Chung, J. H., Kim, J., Kim, K. S., and Han, Y. S. (May, 2017) New role of human ribosomal protein S3: Regulation of cell cycle via phosphorylation by cyclin-dependent kinase 2. Oncol Lett, 13(5), 3681–3687.

77. Russo, A., Catillo, M., Esposito, D., Briata, P., Pietropaolo, C., and Russo, G. (Sep, 2011) Autoregulatory circuit of human rpL3 expression requires hnRNP H1, NPM and KHSRP. Nucleic Acids Res., 39(17), 7576–7585.

78. Takei, S., Togo-Ohno, M., Suzuki, Y., and Kuroyanagi, H. (Mar, 2016) Evolutionarily conserved autoregulation of alternative pre-mRNA splicing by ribosomal protein L10a. Nucleic Acids Res.,.

79. Ruskin, B., Zamore, P. D., and Green, M. R. (Jan, 1988) A factor, U2AF, is required for U2 snRNP binding and splicing complex assembly. Cell, 52(2), 207–219.

80. Wu, S., Romfo, C. M., Nilsen, T. W., and Green, M. R. (Dec, 1999) Functional recognition of the 3’ splice site AG by the splicing factor U2AF35. Nature, 402(6763), 832–835.

81. Berglund, J. A., Abovich, N., and Rosbash, M. (Mar, 1998) A cooperative interaction between U2AF65 and mBBP/SF1 facilitates branchpoint region recognition. Genes Dev., 12(6), 858–867.

82. Jenkins, J. L. and Kielkopf, C. L. (05, 2017) Splicing Factor Mutations in Myelodysplasias: Insights from Spliceosome Structures. Trends Genet., 33(5), 336–348.

83. Kondrashov, F. A. and Koonin, E. V. (Nov, 2001) Origin of alternative splicing by tandem exon duplication. Hum. Mol. Genet., 10(23), 2661–2669.

84. Jourdain, A. A., Popow, J., de la Fuente, M. A., Martinou, J. C., Anderson, P., and Simarro, M. (Nov, 2017) The FASTK family of proteins: emerging regulators of mitochondrial RNA biology. Nucleic Acids Res., 45(19), 10941–10947.

85. Wang, A., Zhao, C., Liu, X., Su, W., Duan, G., Xie, Z., Chu, S., and Gao, Y. (Jan, 2018) Knockdown of TBRG4 affects tumorigenesis in human H1299 lung cancer cells by regulating DDIT3, CAV1 and RRM2. Oncol Lett, 15(1), 121–128.

86. Hermsen, R., Ursem, B., and ten Wolde, P. R. (Jun, 2010) Combinatorial gene regulation using auto-regulation. PLoS Comput. Biol., 6(6), e1000813.

87. Hanamura, A. and Aiba, H. (Aug, 1991) Molecular mechanism of negative autoregulation of Escherichia coli crp gene. Nucleic Acids Res., 19(16), 4413–4419.

88. Bateman, E. (1998) Autoregulation of eukaryotic transcription factors. Prog. Nucleic Acid Res. Mol. Biol., 60, 133–168.

89. Lyabin, D. N., Eliseeva, I. A., Skabkina, O. V., and Ovchinnikov, L. P. (2011) Interplay between Y-box-binding protein 1 (YB-1) and poly(A) binding protein (PABP) in specific regulation of YB-1 mRNA translation. RNA Biol, 8(5), 883–892.

90. Schoenberg, D. R. and Maquat, L. E. (Mar, 2012) Regulation of cytoplasmic mRNA decay. Nat. Rev. Genet., 13(4), 246–259.

91. Wheeler, E. C., Van Nostrand, E. L., and Yeo, G. W. (01, 2018) Advances and challenges in the detection of transcriptome-wide protein-RNA interactions. Wiley Interdiscip Rev RNA, 9(1).

92. Pervouchine, D. D. (Jun, 2018) Towards Long-Range RNA Structure Prediction in Eukaryotic Genes. Genes (Basel), 9(6).

93. Lunde, B. M., Moore, C., and Varani, G. (Jun, 2007) RNA-binding proteins: modular design for efficient function. Nat. Rev. Mol. Cell Biol., 8(6), 479–490.

94. Mitrophanov, A. Y. and Groisman, E. A. (Jun, 2008) Positive feedback in cellular control systems. Bioessays, 30(6), 542–555.

95. Martinez-Pastor, M., Lancaster, W. A., Tonner, P. D., Adams, M. W. W., and Schmid, A. K. (Sep, 2017) A transcription network of interlocking positive feedback loops maintains intracellular iron balance in archaea. Nucleic Acids Res., 45(17), 9990–10001.

96. Ge, Y. and Porse, B. T. (Mar, 2014) The functional consequences of intron retention: alternative splicing coupled to NMD as a regulator of gene expression. Bioessays, 36(3), 236–243.

