## Supplementary material for "Novel autoregulatory cases of alternative splicing coupled with nonsense-mediated mRNA decay"

D. Pervouchine *et al.*

November 8, 2018

S4 GO enrichment analysis (Biological Process) of the 200 genes with the largest fold change between UPF1/XRN1 co-depletion and control. Top 10 GO terms with P-value$< 10^{-3}$ are shown. . 7
S5 GO enrichment analysis (Biological Process) of the genes hosting differentially spliced exons (adjusted p-value$< 0.05$) in UPF1/XRN1 co-depletion and control. The top 15 GO terms with P-value$<10^{-3}$ are shown. . . . 8

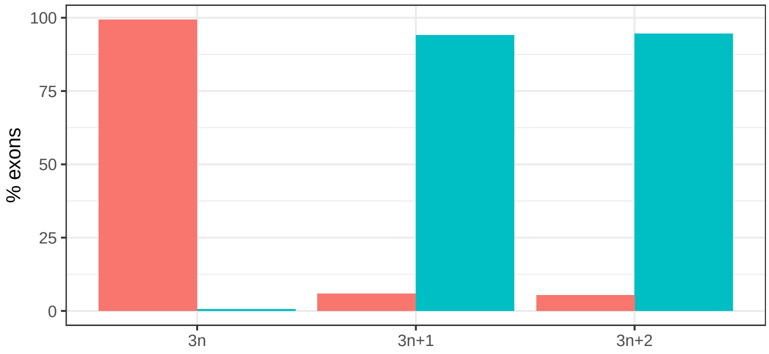

Figure S1: The percentage of essential (green) and non-essential (red) internal exons among all exons annotated in the GENCODE v19 database [1].

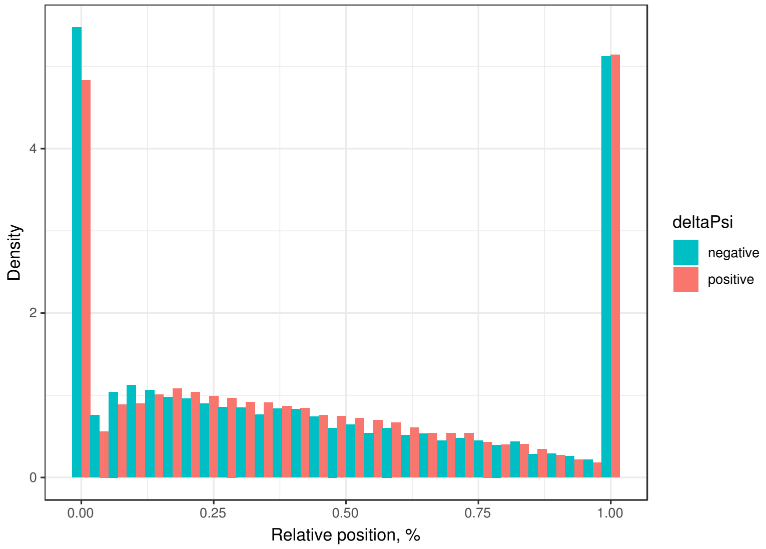

Figure S2: Relative positions within transcript of exons reactive to UPF1/XRN1 co-depletion.

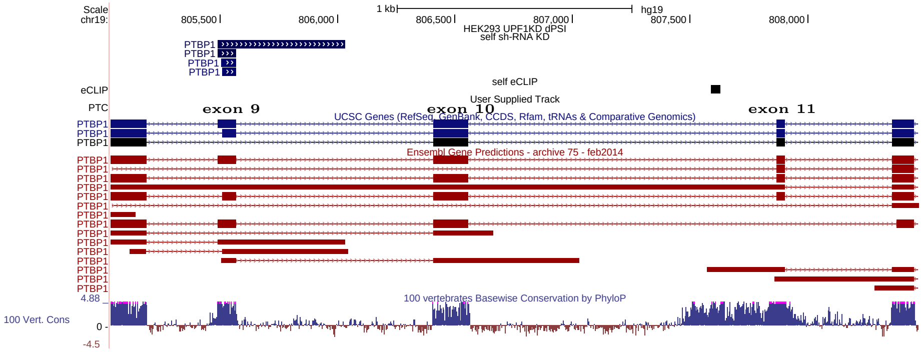

Figure S3: Essential exons in PTBP1 gene. The isoform with exon 11 skipping is not shown in Genome Browser

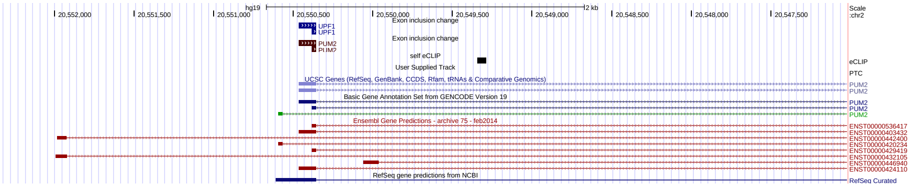

Figure S4: Essential upstream exons in PUM2 gene.

Table S1: The list of RNA-binding proteins, eCLIP, and shRNA-KD followed RNA-seq experiment accession numbers [2, 3]

| RBP | eCLIP | | shRNA KD + RNA-seq | |
|---|---|---|---|---|
| | K562 | HepG2 | K562 | HepG2 |
| AARS | ENCSR825SVO | N/A | ENCSR599UDS | ENCSR547NWD |
| AGGF1 | ENCSR725ARB | ENCSR543TPH | ENCSR812TLY | N/A |
| AKAP8L | ENCSR206RXT | N/A | ENCSR809ISU | ENCSR807ODB |
| BCCIP | N/A | ENCSR485QCG | ENCSR606QIX | ENCSR570CWH |
| BUD13 | ENCSR663WES | ENCSR830BSQ | ENCSR267RHP | ENCSR382QKD |
| CDC40 | N/A | ENCSR815VVI | N/A | N/A |
| CPSF6 | ENCSR532VUB | N/A | ENCSR384BDV | ENCSR676EKU |
| CSTF2 | N/A | ENCSR384MWO | ENCSR885YOI | ENCSR815JDY |
| CSTF2T | ENCSR840DRD | ENCSR919HSE | ENCSR286OKW | ENCSR914WQV |
| DDX24 | ENCSR999WKT | N/A | ENCSR067LLB | ENCSR300IEW |
| DDX3X | ENCSR930BZL | ENCSR648LAH | ENCSR000KYM | ENCSR637JLM |
| DDX42 | ENCSR576SHT | N/A | N/A | N/A |
| DDX55 | ENCSR923NKN | ENCSR845VGB | ENCSR856CJK | ENCSR964YTW |
| DDX59 | N/A | ENCSR214BZA | N/A | ENCSR598GKQ |
| DDX6 | ENCSR893EFU | ENCSR141OIM | ENCSR119QWQ | ENCSR147ZBD |
| DGCR8 | ENCSR947JVR | ENCSR061SZV | N/A | N/A |
| DHX30 | ENCSR529GSJ | ENCSR565DGW | ENCSR345VVZ | ENCSR853PBF |
| DKC1 | N/A | ENCSR301TFY | ENCSR494UDF | ENCSR118KUN |
| DROSHA | ENCSR653HQC | ENCSR834YLD | ENCSR624XHG | N/A |
| EFTUD2 | ENCSR844RVX | ENCSR527DXF | ENCSR117WLY | ENCSR620OKS |
| EIF3D | N/A | ENCSR041NUV | ENCSR660ETT | ENCSR788HVK |
| EIF3H | N/A | ENCSR916XIV | N/A | N/A |
| EIF4G2 | ENCSR307YIW | N/A | ENCSR040FSN | ENCSR152MON |
| EWSR1 | ENCSR887LPK | N/A | ENCSR831YGP | ENCSR532ZPP |
| EXOSC5 | ENCSR013CTQ | N/A | N/A | N/A |
| FAM120A | ENCSR006OEQ | ENCSR987NYS | ENCSR492BKM | ENCSR047VPW |
| FASTKD2 | ENCSR887FHF | ENCSR023UHL | ENCSR608IAI | ENCSR716WZH |
| FKBP4 | N/A | ENCSR018ZUE | ENCSR379VXW | ENCSR639LKS |
| FMR1 | ENCSR331VNX | N/A | ENCSR555LCE | ENCSR905HID |
| FTO | ENCSR989SMC | N/A | ENCSR688GVV | ENCSR389HFU |
| FUBP3 | N/A | ENCSR486YGP | ENCSR373KOF | ENCSR755KOM |
| FXR1 | ENCSR774RFN | N/A | ENCSR780YFF | ENCSR009PPI |
| FXR2 | ENCSR224QWC | N/A | ENCSR577XBW | N/A |
| GEMIN5 | ENCSR238CLX | N/A | ENCSR398GHW | ENCSR771QMJ |
| GNL3 | ENCSR301UQM | N/A | N/A | N/A |
| GPKOW | ENCSR647CLF | N/A | ENCSR967QNT | ENCSR968YWY |
| GRSF1 | N/A | ENCSR668MJX | ENCSR835RMN | ENCSR674KDQ |
| GRWD1 | N/A | ENCSR893NWB | ENCSR528ASX | ENCSR850FEH |
| GTF2F1 | ENCSR736AAG | ENCSR265ZIS | ENCSR188IPO | ENCSR295XKC |
| HLTF | ENCSR589YHM | N/A | ENCSR958NDU | ENCSR010ZMZ |
| HNRNPA1 | ENCSR154HRN | ENCSR769UEW | ENCSR048BWH | ENCSR182DAW |
| HNRNPC | N/A | ENCSR550DVK | ENCSR634KBO | ENCSR052IYH |
| HNRNPK | ENCSR268ETU | ENCSR828ZID | ENCSR529JNJ | ENCSR853ZJS |
| HNRNPM | ENCSR412NOW | ENCSR267UCX | ENCSR746NIM | ENCSR995JMS |
| HNRNPU | ENCSR520BZQ | ENCSR240MVJ | ENCSR047IUS | ENCSR308IKH |

| | | | | |
|---|---|---|---|---|
| HNRNPUL1 | ENCSR571VHI | ENCSR755TJC | ENCSR034VBA | ENCSR689ZJC |
| IGF2BP1 | ENCSR975KIR | ENCSR744GEU | ENCSR629EWX | ENCSR708GKW |
| IGF2BP2 | ENCSR062NNB | N/A | ENCSR952RRH | ENCSR478FJK |
| IGF2BP3 | N/A | ENCSR993OLA | ENCSR302JQA | ENCSR710NWE |
| ILF3 | ENCSR438KWZ | ENCSR786TSC | ENCSR269HQA | ENCSR942MBU |
| KHDRBS1 | ENCSR628IDK | N/A | ENCSR023HWI | ENCSR784FTX |
| KHSRP | ENCSR438GZQ | N/A | ENCSR561CBC | ENCSR850CKU |
| LARP4 | ENCSR888YTT | ENCSR805SRN | ENCSR866XLI | ENCSR744PAQ |
| LARP7 | ENCSR456KXI | ENCSR961OKA | ENCSR770OWW | ENCSR624FBY |
| LIN28B | ENCSR970NKP | ENCSR861GYE | ENCSR598YQX | ENCSR927SLP |
| LSM11 | ENCSR022BVV | ENCSR135VMS | ENCSR762FEO | ENCSR883BXR |
| MATR3 | N/A | ENCSR290VLT | ENCSR792XFP | ENCSR492UFS |
| METAP2 | ENCSR303OQD | N/A | ENCSR952QDQ | ENCSR992JGE |
| MTPAP | ENCSR200DKE | N/A | ENCSR631RFX | ENCSR701GSV |
| NCBP2 | ENCSR484LTQ | ENCSR018RVZ | ENCSR361LBE | ENCSR030ARO |
| NKRF | N/A | ENCSR277DEO | N/A | ENCSR517JDK |
| NOL12 | N/A | ENCSR820DQJ | ENCSR227AVS | ENCSR643UFV |
| NOLC1 | ENCSR001VAC | ENCSR194HZU | N/A | N/A |
| NONO | ENCSR861PAR | N/A | ENCSR398HXV | ENCSR647NYX |
| NPM1 | ENCSR867DSZ | N/A | ENCSR346DZQ | ENCSR016XPB |
| NSUN2 | ENCSR081JYH | N/A | ENCSR829EFL | ENCSR629RUG |
| PCBP2 | N/A | ENCSR339FUY | ENCSR648QFY | N/A |
| POLR2G | N/A | ENCSR820WHR | N/A | N/A |
| PPIG | N/A | ENCSR097NEE | ENCSR529MBZ | ENCSR620HAA |
| PPIL4 | ENCSR197INS | N/A | ENCSR556FNN | ENCSR851KEX |
| PRPF8 | ENCSR534YOI | ENCSR121NVA | ENCSR137HKS | ENCSR998MZP |
| PTBP1 | ENCSR981WKN | ENCSR384KAN | ENCSR527IVX | ENCSR064DXG |
| PUM2 | ENCSR661ICQ | N/A | ENCSR118XYK | ENCSR210DML |
| PUS1 | ENCSR291XPT | N/A | ENCSR618IQH | ENCSR296ERI |
| QKI | ENCSR366YOG | ENCSR570WLM | ENCSR256PLH | ENCSR330YOU |
| RBFOX2 | ENCSR756CKJ | ENCSR987FTF | ENCSR336DFS | ENCSR767LLP |
| RBM15 | ENCSR196INN | ENCSR754NDA | ENCSR385UPQ | ENCSR599PXD |
| RBM22 | ENCSR295OKT | ENCSR456JJQ | ENCSR947OIM | ENCSR330KHN |
| RBM5 | N/A | ENCSR489ABS | N/A | N/A |
| RPS11 | ENCSR269AJF | N/A | N/A | N/A |
| RPS3 | ENCSR120EAR | ENCSR766FAC | N/A | N/A |
| SAFB2 | ENCSR943MHU | N/A | ENCSR770LYW | ENCSR110ZYD |
| SBDS | ENCSR059CWF | N/A | ENCSR219DXZ | ENCSR343DHN |
| SERBP1 | ENCSR121GQH | N/A | ENCSR925RNE | ENCSR820ROH |
| SF3A3 | N/A | ENCSR331MIC | ENCSR454KYR | ENCSR374NMJ |
| SF3B1 | ENCSR133QEA | N/A | ENCSR047QHX | ENCSR896CFV |
| SF3B4 | ENCSR267OLV | ENCSR279UJF | ENCSR081XRA | ENCSR148MQK |
| SFPQ | N/A | ENCSR965DLL | ENCSR535YPK | ENCSR782MXN |
| SLBP | ENCSR483NOP | N/A | ENCSR112YTD | ENCSR519KXM |
| SLTM | N/A | ENCSR351PVI | ENCSR234YMW | ENCSR185JGT |
| SMNDC1 | ENCSR658IQB | ENCSR373ODC | ENCSR408SDL | ENCSR995ZGJ |
| SND1 | ENCSR128VXC | ENCSR061EVO | ENCSR232XRZ | ENCSR398LZW |
| SRSF1 | ENCSR432XUP | ENCSR989VIY | ENCSR066VOO | ENCSR094KBY |
| SRSF7 | ENCSR468FSW | ENCSR513NDD | ENCSR464ADT | ENCSR017PRS |

| | | | | |
|---|---|---|---|---|
| SRSF9 | N/A | ENCSR773KRC | N/A | ENCSR597XHH |
| SUB1 | N/A | ENCSR406OOZ | ENCSR047AJA | ENCSR997FOT |
| SUGP2 | N/A | ENCSR506UPY | ENCSR192BPV | ENCSR837QDN |
| SUPV3L1 | N/A | ENCSR580MFX | ENCSR778SIU | ENCSR995RPB |
| TAF15 | ENCSR568DZW | ENCSR841EQA | ENCSR611ZAL | ENCSR998RZI |
| TARDBP | ENCSR584TCR | N/A | ENCSR134JRE | ENCSR527QNC |
| TBRG4 | ENCSR506OTC | ENCSR916SRV | ENCSR079IPT | ENCSR741YCA |
| TIA1 | ENCSR057DWB | ENCSR623VEQ | ENCSR694LKY | ENCSR057GCF |
| TRA2A | ENCSR365NVO | ENCSR314UMJ | ENCSR916WOI | ENCSR030GZQ |
| TROVE2 | ENCSR539ZTS | ENCSR993FMY | ENCSR060KRD | ENCSR946OFN |
| U2AF1 | ENCSR862QCH | ENCSR328LLU | ENCSR342EDG | ENCSR372UWV |
| U2AF2 | ENCSR893RAV | ENCSR202BFN | ENCSR904CJQ | ENCSR426UUG |
| U2AF2 | ENCSR893RAV | ENCSR202BFN | ENCSR904CJQ | ENCSR622MCX |
| UCHL5 | ENCSR349KMG | ENCSR490IEE | ENCSR678MVE | ENCSR684HTV |
| UPF1 | ENCSR456ASB | N/A | ENCSR251ABP | ENCSR689MIY |
| XPO5 | N/A | ENCSR921SXC | ENCSR453HKS | ENCSR778RWJ |
| XRCC6 | ENCSR258QKO | ENCSR571ROL | ENCSR232CPD | ENCSR500WHE |
| XRN2 | ENCSR657TZB | ENCSR655NZA | ENCSR717SJA | ENCSR347ZHQ |
| YBX3 | ENCSR529FKI | N/A | ENCSR306EIU | ENCSR494VSD |
| YWHAG | ENCSR867ZVK | N/A | N/A | N/A |
| ZC3H11A | ENCSR712IAG | ENCSR907GUB | N/A | N/A |
| ZNF622 | ENCSR657TZZ | N/A | N/A | ENCSR518JXY |
| ZRANB2 | ENCSR663NRA | N/A | ENCSR850PWM | ENCSR081QQH |

Table S2: GO enrichment analysis (Biological Process) of Genes with at least one NMD transcript (GWN) using GOrilla [4].

| GO Term | Description | p-value | FDR q-value |
|---|---|---|---|
| GO:0003824 | catalytic activity | 1.04E-39 | 4.27E-36 |
| GO:0036094 | small molecule binding | 3.91E-23 | 8.01E-20 |
| GO:1901265 | nucleoside phosphate binding | 9.60E-23 | 1.31E-19 |
| GO:0000166 | nucleotide binding | 1.29E-22 | 1.32E-19 |
| GO:0005524 | ATP binding | 3.56E-16 | 2.92E-13 |
| GO:0005488 | binding | 1.22E-15 | 8.34E-13 |
| GO:0035639 | purine ribonucleoside triphosphate binding | 1.50E-15 | 8.81E-13 |
| GO:0003723 | RNA binding | 1.59E-15 | 8.15E-13 |
| GO:0030554 | adenyl nucleotide binding | 1.90E-15 | 8.67E-13 |
| GO:0032559 | adenyl ribonucleotide binding | 2.00E-15 | 8.21E-13 |
| GO:0032549 | ribonucleoside binding | 3.31E-15 | 1.24E-12 |
| GO:0001883 | purine nucleoside binding | 3.69E-15 | 1.26E-12 |
| GO:0032550 | purine ribonucleoside binding | 4.11E-15 | 1.30E-12 |
| GO:0001882 | nucleoside binding | 5.01E-15 | 1.47E-12 |
| GO:0005515 | protein binding | 6.46E-15 | 1.77E-12 |

Table S3: The top 15 genes with the largest gene expression fold change between UPF1/XRN1 co-depletion and control.

| ID | baseMean | log2FoldChange | p-value | Gene |
|---|---|---|---|---|
| TCONS_00166954 | 5365.64 | 2.562 | 0.01388 | BRD2 |
| TCONS_00034807 | 18856.60 | 2.539 | 0.01384 | SNHG1 |
| TCONS_00176132 | 8304.77 | 2.534 | 0.01429 | CCT6P3 |
| TCONS_00176130 | 3356.49 | 2.525 | 0.01538 | CCT6P3 |
| TCONS_00001342 | 7025.46 | 2.505 | 0.01522 | SRRM1 |
| TCONS_00070942 | 3337.18 | 2.505 | 0.01596 | 61E3.4 |
| TCONS_00159029 | 3264.56 | 2.498 | 0.01620 | MATR3 |
| TCONS_00070925 | 2505.97 | 2.493 | 0.01679 | SMG1P1 |
| TCONS_00019843 | 2258.43 | 2.492 | 0.01721 | AGAP6 |
| TCONS_00163703 | 2612.10 | 2.484 | 0.01699 | FAM13B |
| TCONS_00101194 | 4421.11 | 2.478 | 0.01646 | SAE1 |
| TCONS_00182764 | 2072.64 | 2.478 | 0.01765 | DNAJC2 |
| TCONS_00127428 | 9299.07 | 2.478 | 0.01589 | RBM39 |
| TCONS_00010353 | 2930.28 | 2.477 | 0.01796 | CROCCP2 |
| TCONS_00198400 | 2045.45 | 2.477 | 0.01773 | FBXW2 |

Table S4: GO enrichment analysis (Biological Process) of the 200 genes with the largest fold change between UPF1/XRN1 co-depletion and control. Top 10 GO terms with P-value$< 10^{-3}$ are shown.

| GOBPID | ExpCount | Count | Size | Term |
|---|---|---|---|---|
| GO:0006396 | 7 | 37 | 936 | RNA processing |
| GO:0006397 | 3 | 27 | 491 | mRNA processing |
| GO:0008380 | 3 | 23 | 428 | RNA splicing |
| GO:0016071 | 6 | 30 | 812 | mRNA metabolic process |
| GO:0000377 | 2 | 19 | 323 | RNA splicing, via transesterification reactions with bulged adenosine as nucleophile |
| GO:0000398 | 2 | 19 | 323 | mRNA splicing, via spliceosome |
| GO:0000375 | 2 | 19 | 326 | RNA splicing, via transesterification reactions |
| GO:0090304 | 37 | 65 | 5206 | nucleic acid metabolic process |
| GO:0050684 | 1 | 9 | 112 | regulation of mRNA processing |
| GO:0043484 | 0 | 6 | 118 | regulation of RNA splicing |

Table S5: GO enrichment analysis (Biological Process) of the genes hosting differentially spliced exons (adjusted p-value$< 0.05$) in UPF1/XRN1 co-depletion and control. The top 15 GO terms with P-value$< 10^{-3}$ are shown.

| GOBPID | OddsRatio | ExpCount | Count | Size | Term |
|---|---|---|---|---|---|
| GO:0043484 | 30.773 | 0 | 6 | 118 | regulation of RNA splicing |
| GO:0033119 | 103.119 | 0 | 4 | 25 | negative regulation of RNA splicing |
| GO:0006396 | 8.886 | 2 | 12 | 936 | RNA processing |
| GO:0000377 | 15.447 | 1 | 8 | 323 | RNA splicing, via transesterification reactions with bulged adenosine as nucleophile |
| GO:0000398 | 15.447 | 1 | 8 | 323 | mRNA splicing, via spliceosome |
| GO:0000375 | 15.298 | 1 | 8 | 326 | RNA splicing, via transesterification reactions |
| GO:0048024 | 38.162 | 0 | 5 | 78 | regulation of mRNA splicing, via spliceosome |
| GO:0008380 | 11.514 | 1 | 8 | 428 | RNA splicing |
| GO:0050684 | 25.984 | 0 | 5 | 112 | regulation of mRNA processing |
| GO:0016071 | 7.934 | 2 | 10 | 812 | mRNA metabolic process |
| GO:0006397 | 9.975 | 1 | 8 | 491 | mRNA processing |
| GO:0048025 | 92.663 | 0 | 3 | 20 | negative regulation of mRNA splicing, via spliceosome |
| GO:0050686 | 58.310 | 0 | 3 | 30 | negative regulation of mRNA processing |
| GO:0006406 | 20.329 | 0 | 4 | 110 | mRNA export from nucleus |
| GO:0071427 | 20.329 | 0 | 4 | 110 | mRNA-containing ribonucleoprotein complex export from nucleus |

**SupplementaryDataFie 1:** SupplementaryDataFile1.tsv

**SupplementaryDataFie 2:** SupplementaryDataFile2.tsv

### Supplementary References

[1] Church, D. M., Schneider, V. A., Graves, T., Auger, K., Cunningham, F., Bouk, N., Chen, H. C., Agarwala, R., McLaren, W. M., Ritchie, G. R., Albracht, D., Kremitzki, M., Rock, S., Kotkiewicz, H., Kremitzki, C., Wollam, A., Trani, L., Fulton, L., Fulton, R., Matthews, L., Whitehead, S., Chow, W., Torrance, J., Dunn, M., Harden, G., Threadgold, G., Wood, J., Collins, J., Heath, P., Griffiths, G., Pelan, S., Grafham, D., Eichler, E. E., Weinstock, G., Mardis, E. R., Wilson, R. K., Howe, K., Flicek, P., and Hubbard, T. (Jul, 2011) Modernizing reference genome assemblies. *PLoS Biol.,* **9**(7), e1001091.

[2] Dunham, I., Kundaje, A., Aldred, S. F., Collins, P. J., Davis, C. A., Doyle, F., Epstein, C. B., Frietze, S., Harrow, J., Kaul, R., Khatun, J., Lajoie, B. R., Landt, S. G., Lee, B. K., Pauli, F., Rosenbloom, K. R., Sabo, P., Safi, A., Sanyal, A., Shoresh, N., Simon, J. M., Song, L., Trinklein, N. D., Altshuler, R. C., Birney, E., Brown, J. B., Cheng, C., Djebali, S., Dong, X., Dunham, I., Ernst, J., Furey, T. S., Gerstein, M., Giardine, B., Greven, M., Hardison, R. C., Harris, R. S., Herrero, J., Hoffman, M. M., Iyer, S., Kellis, M., Khatun, J., Kheradpour, P., Kundaje, A., Lassmann, T., Li, Q., Lin, X., Marinov, G. K., Merkel, A., Mortazavi, A., Parker, S. C., Reddy, T. E., Rozowsky, J., Schlesinger, F., Thurman, R. E., Wang, J., Ward, L. D., Whitfield, T. W., Wilder, S. P., Wu, W., Xi, H. S., Yip, K. Y., Zhuang, J., Pazin, M. J., Lowdon, R. F., Dillon, L. A., Adams, L. B., Kelly, C. J., Zhang, J., Wexler, J. R., Green, E. D., Good, P. J., Feingold, E. A., Bernstein, B. E., Birney, E., Crawford, G. E., Dekker, J., Elnitski, L., Farnham, P. J., Gerstein, M., Giddings, M. C., Gingeras, T. R., Green, E. D., Guigo, R., Hardison, R. C., Hubbard, T. J., Kellis, M., Kent, W., Lieb, J. D., Margulies, E. H., Myers, R. M., Snyder, M., Stamatoyannopoulos, J. A., Tenenbaum, S. A., Weng, Z., White, K. P., Wold, B., Khatun, J., Yu, Y., Wrobel, J., Risk, B. A., Gunawardena, H. P., Kuiper, H. C., Maier, C. W., Xie, L., Chen, X., Giddings, M. C., Bernstein, B. E., Epstein, C. B., Shoresh, N., Ernst, J., Kheradpour, P., Mikkelsen, T. S., Gillespie, S., Goren, A., Ram, O., Zhang, X., Wang, L., Issner, R., Coyne, M. J., Durham, T., Ku, M., Truong, T., Ward, L. D., Altshuler, R. C., Eaton, M. L., Kellis, M., Djebali, S., Davis, C. A., Merkel, A., Dobin, A., Lassmann, T., Mortazavi, A., Tanzer, A., Lagarde, J., Lin, W., Schlesinger, F., Xue, C., Marinov, G. K., Khatun, J., Williams, B. A., Zaleski, C., Rozowsky, J., Roder, M., Kokocinski, F., Abdelhamid, R. F., Alioto, T., Antoshechkin, I., Baer, M. T., Batut, P., Bell, I., Bell, K., Chakrabortty, S., Chen, X., Chrast, J., Curado, J., Derrien, T., Drenkow, J., Dumais, E., Dumais, J., Duttagupta, R., Fastuca, M., Fejes-Toth, K., Ferreira, P., Foissac, S., Fullwood, M. J., Gao, H., Gonzalez, D., Gordon, A., Gunawardena, H. P., Howald, C., Jha, S., Johnson, R., Kapranov, P., King, B., Kingswood, C., Li, G., Luo, O. J., Park, E., Preall, J. B., Presaud, K., Ribeca, P., Risk, B. A., Robyr, D., Ruan, X., Sammeth, M., Sandhu, K. S., Schaeffer, L., See, L. H., Shahab, A., Skancke, J., Suzuki, A. M., Takahashi, H., Tilgner, H., Trout, D., Walters, N., Wang, H., Wrobel, J., Yu, Y., Hayashizaki, Y., Harrow, J., Gerstein, M., Hubbard, T. J., Reymond, A., Antonarakis, S. E., Hannon, G. J., Giddings, M. C., Ruan, Y., Wold, B., Carninci, P., Guigo, R., Gingeras, T. R., Rosenbloom,

K. R., Sloan, C. A., Learned, K., Malladi, V. S., Wong, M. C., Barber, G. P., Cline, M. S., Dreszer, T. R., Heitner, S. G., Karolchik, D., Kent, W., Kirkup, V. M., Meyer, L. R., Long, J. C., Maddren, M., Raney, B. J., Furey, T. S., Song, L., Grasfeder, L. L., Giresi, P. G., Lee, B. K., Battenhouse, A., Sheffield, N. C., Simon, J. M., Showers, K. A., Safi, A., London, D., Bhinge, A. A., Shestak, C., Schaner, M. R., Kim, S. K., Zhang, Z. Z., Mieczkowski, P. A., Mieczkowska, J. O., Liu, Z., McDaniell, R. M., Ni, Y., Rashid, N. U., Kim, M. J., Adar, S., Zhang, Z., Wang, T., Winter, D., Keefe, D., Birney, E., Iyer, V. R., Lieb, J. D., Crawford, G. E., Li, G., Sandhu, K. S., Zheng, M., Wang, P., Luo, O. J., Shahab, A., Fullwood, M. J., Ruan, X., Ruan, Y., Myers, R. M., Pauli, F., Williams, B. A., Gertz, J., Marinov, G. K., Reddy, T. E., Vielmetter, J., Partridge, E., Trout, D., Varley, K. E., Gasper, C., Bansal, A., Pepke, S., Jain, P., Amrhein, H., Bowling, K. M., Anaya, M., Cross, M. K., King, B., Muratet, M. A., Antoshechkin, I., Newberry, K. M., McCue, K., Nesmith, A. S., Fisher-Aylor, K. I., Pusey, B., DeSalvo, G., Parker, S. L., Balasubramanian, S., Davis, N. S., Meadows, S. K., Eggleston, T., Gunter, C., Newberry, J., Levy, S. E., Absher, D. M., Mortazavi, A., Wong, W. H., Wold, B., Blow, M. J., Visel, A., Pennachio, L. A., Elnitski, L., Margulies, E. H., Parker, S. C., Petrykowska, H. M., Abyzov, A., Aken, B., Barrell, D., Barson, G., Berry, A., Bignell, A., Boychenko, V., Bussotti, G., Chrast, J., Davidson, C., Derrien, T., Despacio-Reyes, G., Diekhans, M., Ezkurdia, I., Frankish, A., Gilbert, J., Gonzalez, J. M., Griffiths, E., Harte, R., Hendrix, D. A., Howald, C., Hunt, T., Jungreis, I., Kay, M., Khurana, E., Kokocinski, F., Leng, J., Lin, M. F., Loveland, J., Lu, Z., Manthravadi, D., Mariotti, M., Mudge, J., Mukherjee, G., Notredame, C., Pei, B., Rodriguez, J. M., Saunders, G., Sboner, A., Searle, S., Sisu, C., Snow, C., Steward, C., Tanzer, A., Tapanari, E., Tress, M. L., van Baren, M. J., Walters, N., Washietl, S., Wilming, L., Zadissa, A., Zhang, Z., Brent, M., Haussler, D., Kellis, M., Valencia, A., Gerstein, M., Reymond, A., Guigo, R., Harrow, J., Hubbard, T. J., Landt, S. G., Frietze, S., Abyzov, A., Addleman, N., Alexander, R. P., Auerbach, R. K., Balasubramanian, S., Bettinger, K., Bhardwaj, N., Boyle, A. P., Cao, A. R., Cayting, P., Charos, A., Cheng, Y., Cheng, C., Eastman, C., Euskirchen, G., Fleming, J. D., Grubert, F., Habegger, L., Hariharan, M., Harmanci, A., Iyengar, S., Jin, V. X., Karczewski, K. J., Kasowski, M., Lacroute, P., Lam, H., Lamarre-Vincent, N., Leng, J., Lian, J., Lindahl-Allen, M., Min, R., Miotto, B., Monahan, H., Moqtaderi, Z., Mu, X. J., O'Geen, H., Ouyang, Z., Patacsil, D., Pei, B., Raha, D., Ramirez, L., Reed, B., Rozowsky, J., Sboner, A., Shi, M., Sisu, C., Slifer, T., Witt, H., Wu, L., Xu, X., Yan, K. K., Yang, X., Yip, K. Y., Zhang, Z., Struhl, K., Weissman, S. M., Gerstein, M., Farnham, P. J., Snyder, M., Tenenbaum, S. A., Penalva, L. O., Doyle, F., Karmakar, S., Landt, S. G., Bhanvadia, R. R., Choudhury, A., Domanus, M., Ma, L., Moran, J., Patacsil, D., Slifer, T., Victorsen, A., Yang, X., Snyder, M., Auer, T., Centanin, L., Eichenlaub, M., Gruhl, F., Heermann, S., Hoeckendorf, B., Inoue, D., Kellner, T., Kirchmaier, S., Mueller, C., Reinhardt, R., Schertel, L., Schneider, S., Sinn, R., Wittbrodt, B., Wittbrodt, J., Weng, Z., Whitfield, T. W., Wang, J., Collins, P. J., Aldred, S. F., Trinklein, N. D., Partridge, E. C., Myers, R. M., Dekker, J., Jain, G., Lajoie, B. R., Sanyal, A., Balasundaram, G., Bates, D. L., Byron, R., Canfield, T. K., Diegel, M. J., Dunn, D., Ebersol, A. K., Frum, T., Garg, K., Gist, E., Hansen, R., Boatman, L., Haugen, E., Humbert, R., Jain,

G., Johnson, A. K., Johnson, E. M., Kutyavin, T. V., Lajoie, B. R., Lee, K., Lotakis, D., Maurano, M. T., Neph, S. J., Neri, F. V., Nguyen, E. D., Qu, H., Reynolds, A. P., Roach, V., Rynes, E., Sabo, P., Sanchez, M. E., Sandstrom, R. S., Sanyal, A., Shafer, A. O., Stergachis, A. B., Thomas, S., Thurman, R. E., Vernot, B., Vierstra, J., Vong, S., Wang, H., Weaver, M. A., Yan, Y., Zhang, M., Akey, J. M., Bender, M., Dorschner, M. O., Groudine, M., MacCoss, M. J., Navas, P., Stamatoyannopoulos, G., Kaul, R., Dekker, J., Stamatoyannopoulos, J. A., Dunham, I., Beal, K., Brazma, A., Flicek, P., Herrero, J., Johnson, N., Keefe, D., Lukk, M., Luscombe, N. M., Sobral, D., Vaquerizas, J. M., Wilder, S. P., Batzoglou, S., Sidow, A., Hussami, N., Kyriazopoulou-Panagiotopoulou, S., Libbrecht, M. W., Schaub, M. A., Kundaje, A., Hardison, R. C., Miller, W., Giardine, B., Harris, R. S., Wu, W., Bickel, P. J., Banfai, B., Boley, N. P., Brown, J. B., Huang, H., Li, Q., Li, J. J., Noble, W. S., Bilmes, J. A., Buske, O. J., Hoffman, M. M., Sahu, A. D., Kharchenko, P. V., Park, P. J., Baker, D., Taylor, J., Weng, Z., Iyer, S., Dong, X., Greven, M., Lin, X., Wang, J., Xi, H. S., Zhuang, J., Gerstein, M., Alexander, R. P., Balasubramanian, S., Cheng, C., Harmanci, A., Lochovsky, L., Min, R., Mu, X. J., Rozowsky, J., Yan, K. K., Yip, K. Y., and Birney, E. (Sep, 2012) An integrated encyclopedia of DNA elements in the human genome. *Nature,* **489**(7414), 57–74.

[3] Sloan, C. A., Chan, E. T., Davidson, J. M., Malladi, V. S., Strattan, J. S., Hitz, B. C., Gabdank, I., Narayanan, A. K., Ho, M., Lee, B. T., Rowe, L. D., Dreszer, T. R., Roe, G., Podduturi, N. R., Tanaka, F., Hong, E. L., and Cherry, J. M. (Jan, 2016) ENCODE data at the ENCODE portal. *Nucleic Acids Res.,* **44**(D1), D726–732.

[4] Eden, E., Navon, R., Steinfeld, I., Lipson, D., and Yakhini, Z. (Feb, 2009) GOrilla: a tool for discovery and visualization of enriched GO terms in ranked gene lists. *BMC Bioinformatics,* **10**, 48.
